# Circuit-motivated generalized affine models characterize stimulus-dependent visual cortical shared variability

**DOI:** 10.1101/2023.12.19.572428

**Authors:** Ji Xia, Anna Jasper, Adam Kohn, Kenneth D. Miller

## Abstract

Correlated variability in the visual cortex is modulated by stimulus properties. The stimulus dependence of correlated variability impacts stimulus coding and is indicative of circuit structure. An affine model combining a factor proportional to mean stimulus response and an additive offset has been proposed to explain how correlated variability in primary visual cortex (V1) depends on stimulus orientations. However, whether the affine model could be extended to explain modulations by other stimulus variables or variability shared between two brain areas is unknown. Motivated by a simple neural circuit mechanism, we modified the affine model to better explain the contrast-dependence of neural variability shared within either primary or secondary visual cortex (V1 or V2) as well as the orientation-dependence of neural variability shared between V1 and V2. Our results bridge neural circuit mechanisms and statistical models, and provide a parsimonious explanation for the stimulus-dependence of correlated variability within and between visual areas.

## Introduction

Neural responses in the visual cortex to repeated presentations of a fixed visual stimulus exhibit trial-to-trial variability. These trial-to-trial fluctuations are correlated or shared among neurons in a local population. There are three primary motivations for investigating and characterizing this shared variability. First, it plays a pivotal role in shaping stimulus coding (Abbott and Dayan, 1999; Averbeck, Latham, and Pouget, 2006; Moreno-Bote et al., 2014; Kohn, Coen-Cagli, et al., 2016). For example, shared variability patterns that steer the population response from one stimulus towards the response to a nearby stimulus can hinder accurate stimulus coding (Moreno-Bote et al., 2014). Second, accumulating evidence suggests that the standard deviation of shared variability at each neuron is modulated by stimuli (Kohn and Smith, 2005; Ponce-Alvarez et al., 2013; Lin et al., 2015; Zylberberg et al., 2016), as well as by behavioral (Stringer et al., 2019; Musall et al., 2019) and task-related (Cohen and Maunsell, 2009; Ruff and Cohen, 2014; Bondy, Haefner, and Cumming, 2018) variables. Consequently, a prevailing argument posits that shared variability does not just reflect stochasticity; instead, it serves to encode additional, complementary information that is absent from the trial-averaged responses and may have computational utility (Haimerl, Savin, and E. Simoncelli, 2019; Echeveste et al., 2020; Lange and Haefner, 2022). Third, the pattern of shared variability reflects the underlying connectivity among neurons (Trousdale et al., 2012; Ocker et al., 2017). Therefore, characterizing shared variability serves as a valuable approach for inferring circuit structure, especially because performing experimental measurements of single neuron input-output function or connectivity are often challenging (Priebe et al., 2004; Hofer et al., 2011; Ko et al., 2011).

In this work, we focus on investigating and characterizing how shared variability depends on stimuli. A comprehensive understanding of the stimulus-dependent nature of shared variability lays the groundwork for quantifying its impact on stimulus coding and inferring circuit structure.

Three models of stimulus-modulated shared variability have been considered in most previous work (Goris, Movshon, and E. P. Simoncelli, 2014; Rabinowitz et al., 2015; Lin et al., 2015; Arandia-Romero et al., 2016): the amplitude (standard deviation) of a given neuron’s shared variability might be multiplicative (proportional to the neuron’s stimulus response), additive (stimulus independent), or affine (combination of a multiplicative and an additive component). For neurons in the visual cortex, previous work found that multiplicative (Goris, Movshon, and E. P. Simoncelli, 2014) or affine (Lin et al., 2015; Arandia-Romero et al., 2016) models outperform the additive model in explaining the neural data. However, further work is needed to thoroughly characterize stimulus-dependent shared variability. It is possible that the underlying modulation takes other more complicated forms than the three proposed models. Furthermore, these models cannot account for the fact that shared variability is suppressed with increasing stimulus contrast (M. M. Churchland et al., 2010; Hennequin et al., 2018). Moreover, these models are statistical in nature and are lack a foundation in neural circuitry mechanism. In this study, we will demonstrate that consideration of such mechanisms suggests alternative forms for the modulation of shared variability.

Trial-to-trial variability is not only correlated across neurons within a brain area, but also across neurons from two connected brain areas (Nowak et al., 1999; Jia, Tanabe, and Kohn, 2013; Pooresmaeili, Poort, and Roelfsema, 2014; Ruff and Cohen, 2016a; Semedo, Zandvakili, et al., 2019; Ruff and Cohen, 2019). This correlation of neural activity between brain areas is often interpreted as area-to-area communication. Characterizing the shared variability between brain areas is vital for understanding its impact on joint stimulus coding by multiple brain areas and for inferring inter-area connectivity. Recent studies found that variability shared between areas has lower dimensionality compared with that shared within each area (Semedo, Zandvakili, et al., 2019), which suggests flexible inter-areal communication occurs through a low-dimensional subspace. However, how this inter-areal shared variability is modulated by the external stimuli, and in particular whether this modulation can also be captured by the previously proposed multiplicative or affine statistical models, remains unknown (Ruff and Cohen, 2016b; Kohn, Jasper, et al., 2020).

Here, we investigated how shared variability within and between visual areas depends on the stimulus. We analyzed electrophysiology data simultaneously recorded in monkeys from areas V1 and V2 in response to visual stimulation of drifting gratings with different orientations and contrasts. We studied a simple circuit mechanism for explaining the previously observed affine-like shared variability in V1 (Lin et al., 2015; Arandia-Romero et al., 2016). We assumed that neural variability is the local fluctuation around a steady state of the recurrent circuit, driven by stimulus-independent external noise. Based on this mechanism, we made three predictions: 1) When firing rates are low, affine-like shared variability should be commonly observed in different brain areas with rectified power law activation functions. 2) In V1, if we consider varying both contrasts and orientations, affine models must be modified to allow contrast-specific coefficients, due to suppression of shared variability with increasing contrast by recurrent dynamics (Hennequin et al., 2018). 3) With minimal assumptions, variability shared between V1 and V2 should also be affine-like across stimulus orientations. Consistent with 1), we found that affine models also explain V2 data well when we varied the stimulus orientations. Consistent with 2), when we varied both contrasts and orientations, we found that a generalized affine model with contrast-specific coefficients best explained shared variability in V2. Consistent with 3), affine models parsimoniously explained how the inter-areal variability shared between V1 and V2 depends on stimulus orientations. We also considered a generalized model that can have an arbitrary form of shared variability for each stimulus. This model slightly outperformed affine models when we had sufficient data to fit it, but differences were small.

Our work extends previous work in characterizing the nature of shared variability within a brain area to shared variability between brain areas. Moreover, we provide a mechanistic explanation for the commonly observed affine-like shared variability. Importantly, we propose an alternative form of stimulus-dependent shared variability that well-explains the data when varying both orientation and contrast.

## Results

### Box 1

**How does neural variability depend on the stimulus in a recurrent neural network?**

In this box, we show that in a recurrent neural network, one should expect neural variability (fluctuation around the fixed point) to depend on the stimulus (external input), if we have a nonlinear activation function for the neurons.

Let’s consider neural dynamics governed by a standard rate network equation:

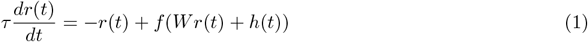

Here, *τ* is the time constant of neurons. *r*(*t*) is a vector that denotes the firing rates of all the neurons at time *t. W* is the matrix of recurrent connectivity between neurons. *h*(*t*) is a vector that denotes the external input received by all the neurons at time *t. f* (*·*) is the activation function, which is applied element-by-element to its vector argument.

Consider the case where *h*(*t*) = *h*^*^ is a constant over time. Then *r*(*t*) will evolve to a fixed point *r*^*^ that satisfies

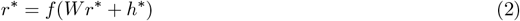

provided that this fixed point is stable. We are interested in how *r*(*t*) fluctuates around *r*^*^, if *h*(*t*) = *h*^*^ + *δh*(*t*), where *δh*(*t*) denotes some external input noise. In other words, we want to know *δr* that satisfies the following equation:

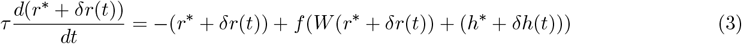

As in (Trousdale et al., 2012; Ocker et al., 2017; Hennequin et al., 2018), we assume that both *δr* and *δh* are sufficiently small that we can Taylor expand the second term on the right hand side of eq. 3 around the fixed point and only keep the first-order term:

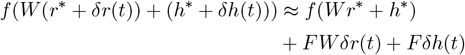

*F* is a diagonal matrix with diagonal entries *f*^*′*^(*Wr*^*^ + *h*^*^), which is the vector of neuronal gains – the slope of the activation function at each neuron’s fixed-point firing rate.

With eq. 2, we can simplify eq. 3 to get:

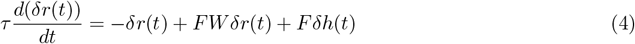

If we further assume that the external input noise *δh*(*t*) varies slowly over time (much slower than *τ*), the left-hand side of eq. 4 will tend to zero, thus we can get an expression for *δr*(*t*):

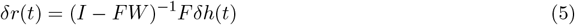

If the activation function *f* (*·*) is nonlinear, then *F* changes with neuronal activation level and thus depends on *r*^*^, which, in turn, depends on the stimulus *h*^*^. Consequently, *δr*(*t*) also depends on the stimulus through *F*, as discussed before in Doiron et al., 2016.

To gain more intuition on what *δr*(*t*) looks like, consider a special case where *f* (*x*) = *kx*^*n*^, then we have *F* = diag(*nk*^1*/n*^*r*^*(1−1*/n*)^). As a motivation for the choice, it has been shown that this power-law nonlinearity well describes the responses of visual cortical neurons (Miller and Troyer, 2002; Hansel and Vreeswijk, 2002; Fourcaud-Trocmé et al., 2003), with evidence from in-vivo whole cell recordings in V1 (Priebe et al., 2004).

If all the eigenvalues of *FW* have absolute values smaller than 1, then we can expand (*I* − *FW*)^−1^ to obtain

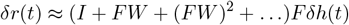

The largest (first) term is *Fδh*(*t*) ∝ diag(*r*^*(1−1*/n*)^)*δh*(*t*). In this case, *δr*(*t*) tends to be higher for those neurons with higher mean firing rate *r*^*^ under a given stimulus *h*^*^.

If all the eigenvalues of *FW* have absolute values larger than 1, then we can expand (*I* − *FW*)^−1^ = (*I* − (*FW*)^−1^)^−1^(−*FW*)^−1^ to obtain

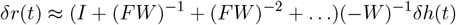

The largest (first) term is −*W*^−1^*δh*(*t*), which is independent of the mean firing rate *r*^*^ under a given stimulus. The higher order terms will be stimulus-dependent through *F*, however, the amplitude of these terms will decrease with increasing *r*^*^. Thus, the stimulus dependence of *δr* becomes weaker as population mean firing rates increase.

In summary, when the mean firing rates *r*^*^ are small enough so that *FW* has a spectral radius less than 1, the amplitude of neural variability *δr* is stimulus-dependent and increases with increasing *r*^*^; when the mean firing rates *r*^*^ are large enough so that all the eigenvalues of *FW* have absolute values larger than 1, the stimulus-dependence of neural variability *δr* becomes weaker with increasing *r*^*^.

### Circuit mechanism for stimulus-dependent shared variability

Numerous works have shown that in the visual cortex, shared variability is low dimensional, such that a large portion of the variance can be explained with a 1-dimensional component or a common fluctuation across neurons (Ecker et al., 2014; Goris, Movshon, and E. P. Simoncelli, 2014; Lin et al., 2015; Arandia-Romero et al., 2016; Huang et al., 2019). To date, there are two hypothesized circuit mechanisms for explaining low-dimensional shared variability: first, it may be inherited from low-dimensional external input noise, and modified by recurrent processing (De La Rocha et al., 2007; Hennequin et al., 2018); second, it may be intrinsically generated by chaotic dynamics in the recurrent circuit (Sompolinsky, Crisanti, and Sommers, 1988; Litwin-Kumar and Doiron, 2012; Huang et al., 2019). In our work, we focused on the first possibility, because this framework is more theoretically tractable and equally biologically plausible, compared to the second possibility.

To investigate the stimulus-dependence of shared variability, we started with deriving an analytical expression for neural variability in the recurrent neural circuit, assuming that external input noise is slow and small (box 1). For the sake of simplicity, we assume for now that external input noise is 1-dimensional, i.e., perfectly correlated across all recipient neurons in our circuit model. Consequently, in this model, neural variability is the same as shared variability, and private variability is not considered. As shown by eq. 5, even if the external input noise was stimulus-independent, the shared variability would depend on the stimulus, because it depends on the neuronal gains (the slopes of the activation function at each neuron’s firing rate), which in turn depend on the stimulus-driven neuronal firing rates, given that the activation function of each neuron is nonlinear.

When would the stimulus-dependence of shared variability be affine? If we further assume that the activation function follows a rectified power law and the firing rates are low enough so that effective recurrent connection (the product of synaptic strengths and neuronal gains) are negligible (see box 1), then the amplitude of shared variability would be proportional to *r*^*(1−1*/n*)^ (Here *r*^*^ is the trial-averaged activity, n is the exponent of the power law in the activation function). The tuning curve of *δr* in practice can be well-approximated by an affine transformation of the tuning curve of *r*^*^. This approximation holds since the exponent 1 − 1*/n* primarily acts to flatten the tuning curve.

To investigate the case where effective recurrent connections are strong, we directly simulated a recurrent neural network with bump-shaped orientation tuning curves, using a classical ring architecture (Rubin, Van Hooser, and Miller, 2015; Hennequin et al., 2018) (Fig. 1A). Neurons in the network receive two external inputs 1) a tuned static input representing the visual stimulation and 2) perfectly correlated external noise that is slow and small (Fig. 1B). We calculated the amplitude of shared variability numerically from the network simulation (see Methods, Fig. 1B) and analytically according to eq. 5 (Fig. 1C).

**Figure 1:**
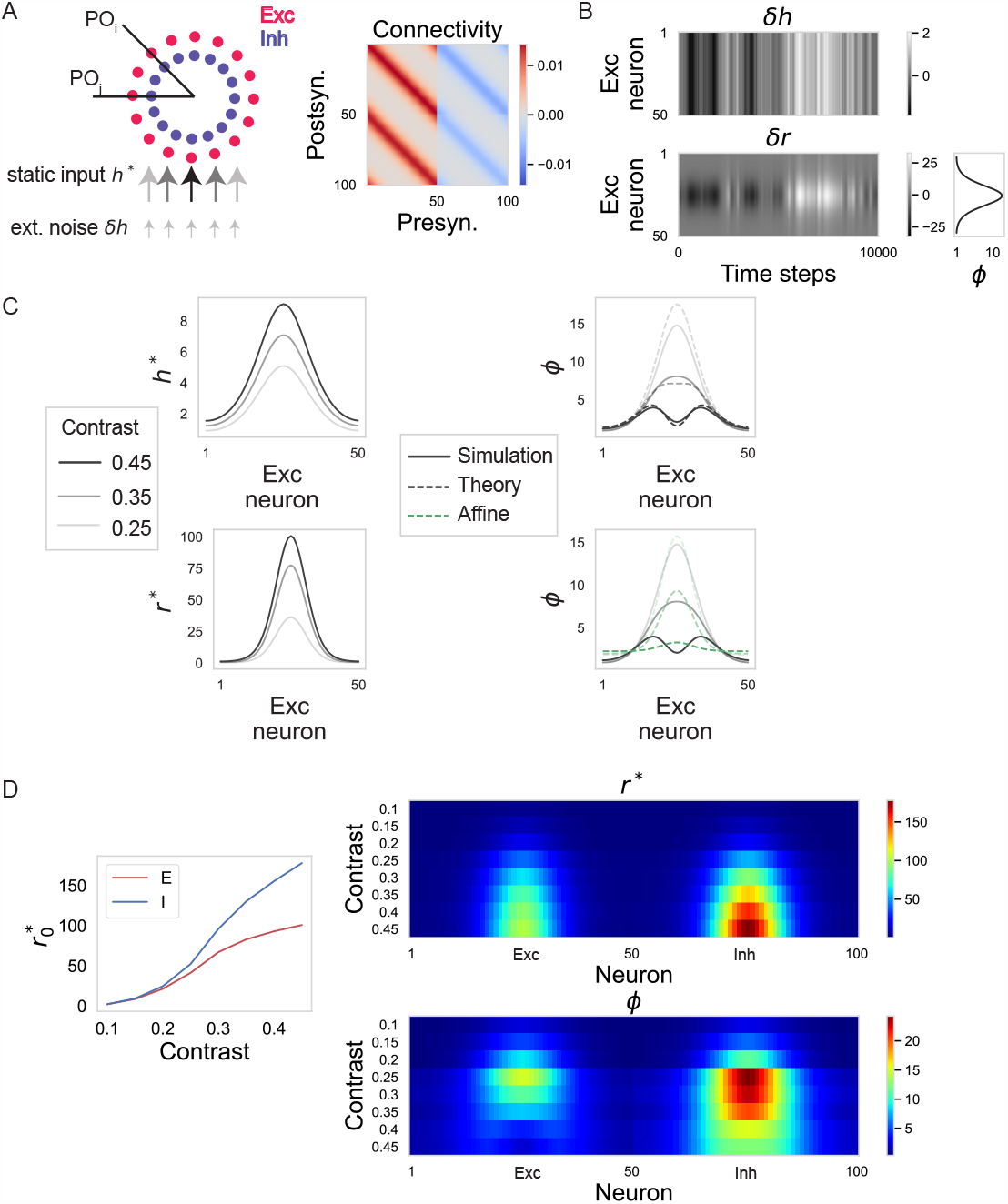
Circuit mechanism for stimulus-dependent shared variability. A. Left: a schematic of the E-I neural circuit with a ring architecture. Excitatory neurons (red circles) and inhibitory neurons (blue circles) have preferred orientations (PO) based on their angular positions on the ring. Two external inputs are denoted by the arrows in the bottom. Right: the connectivity strengths between model neurons. The first 50 neurons are excitatory, the last 50 neurons are inhibitory. For simplicity, we ignore differences in numbers of E vs. I neurons. B. Top: perfectly correlated external noise. Bottom: the corresponding residual responses (responses with time-averaged activity subtracted) given a visual stimulus centered between neurons 25 and 26. On the right, we plot the amplitude of shared variability *ϕ* across neurons. Here we only show results for excitatory neurons. C. Left top: three external static inputs *h*^*^ representing visual inputs with different contrasts received by the model neurons (gray to black denotes low to high contrast). Left bottom: the corresponding time-averaged responses *r*^*^ of the model neurons. Right top: *ϕ* at each neuron from numerical simulation (solid line) and analytical estimation (dashed line; see eq. 5). Right bottom: the prediction of *ϕ* from the affine models (green dashed line) plotted against the simulated *ϕ*. D. Left: The simulated time-averaged firing rates of E and I neurons with PO matching the stimulus orientation, plotted against stimulus contrast. Right: Top: The simulated time-averaged firing rates of all neurons under stimuli with different contrasts. Bottom: The shared variability amplitudes of all neurons under stimuli with different contrast levels.

For simplicity, we set the amplitudes of external noise to be the same across all recipient neurons, so that due to the ring symmetry, the dependence of shared variability on stimulus orientation is the same as its dependence on neuronal preferred orientation (see Supplementary Fig. S1 for similar results when we have external noise with heterogeneous amplitudes across recipient neurons). Interestingly, even when effective recurrent connections are not negligibly small as assumed in box 1, shared variability showed affine-like modulation across orientations for a wide range of contrast levels (Fig. 1C, D). However, the circuit model results also showed two signatures that can’t be captured by affine models, due to recurrent dynamics. First, at high contrast level, the shared variability amplitude exhibits an “M”-shaped dependence on stimulus orientation (Fig 1C); experimentally, “M”-shaped noise correlation has been observed in monkey area MT (Ponce-Alvarez et al., 2013). Second, with increasing contrast, shared variability amplitude first increases and then decreases (Fig. 1D; (Hennequin et al., 2018)); such a decrease in shared variability amplitude with increasing stimulus strength has been observed in many cortical systems (M. M. Churchland et al., 2010). In the supplementary material (S1-S3), we further analyze the behavior of shared variability in the ring circuit model to obtain an intuition for the origin of these two discrepancies from the affine models (building on Hennequin et al., 2018).

In summary, through an analysis of the dynamics in a recurrent network subject to common input noise, we suggest an origin for affine-like shared variability across orientations when the activation function adheres to a rectified power law. Additionally, we establish the need for the development of a novel statistical model to more effectively capture the stimulus-dependent nature of shared variability when both stimulus orientation and contrast are varied.

### Affine models parsimoniously explain shared variability within V1 and V2 across stimulus orientations

To characterize the nature of shared variability in V1 and V2, we first analyzed electrophysiological recordings from macaque V1 and V2 under stimulation by drifting gratings of varying orientations and a single (100%) contrast (see descriptions of dataset 1 in Methods). A given experimental session had 400 trials of each stimulus. Our goal is to identify a statistical model that explains the shared variability across stimuli parsimoniously. We modeled single-trial neural responses as a sum of three elements (Fig. 2A): trial-averaged responses; neural variability shared across neurons, which itself may have multiple components; and neural variability private to each neuron. For neuron *c* on trial *i* with stimulus *s*, its response is modeled as:

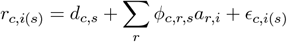

*d*_*c,s*_ is the trial-averaged response of neuron *c* to stimulus *s*; *a*_*r,i*_ is a standard Gaussian variable, *a*_*r,i*_ ∼ 𝒩 (0, 1), that governs how the *r*^*th*^ component of the shared variability varies across trials; *ϕ*_*c,r,s*_ denotes the amplitude of the *r*^*th*^ component of the shared variability for neuron *c* with stimulus *s*; *ϵ*_*c,i*(*s*)_ represents the private variability, which on each trial is a sample from a Gaussian distribution with neuron- and stimulus-specific variance 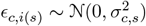.

We considered four types of models with different constraints on the stimulus-dependence of *ϕ* (Fig. 2C). For the additive model, the amplitude of shared variability doesn’t vary across stimuli (*ϕ*_*c,r,s*_ = *ϕ*_*c,r*_). For the multiplicative model, the amplitude of shared variability is proportional to the trial-averaged activity (*ϕ*_*c,r,s*_ = *α*_*c,r*_*d*_*c,s*_). For the affine model, the amplitude of shared variability is constrained to be an affine transformation of the trial-averaged response (*ϕ*_*c,r,s*_ = *α*_*c,r*_*d*_*c,s*_ + *β*_*c,r*_). For the generalized model, there are no constraints on the stimulus-dependence of *ϕ*_*c,r,s*_, as it can vary across *s* freely. In terms of number of parameters in the model, additive model (2*NS* + *NR* parameters) = multiplicative model (2*NS* + *NR* parameters) *<* affine model (2*NS* + 2*NR* parameters) *<* generalized model (2*NS* + *NRS* parameters), where *N* is the number of neurons, *S* is the number of stimuli, and *R* is the number of components of shared variability. If we have enough data so that none of the statistical models overfit, then we should always expect to see that the generalized model performs the best (see Supplementary Fig. S3 for fitting performances of statistical models on the surrogate data with known ground truth). Note that our proposed statistical models have a close relation with factor analysis: the generalized model is equivalent to applying factor analysis separately to the residual responses to each stimulus; the additive model is similar to applying factor analysis to the set of residual responses to all stimuli, except that it assumes stimulus-dependent private variability instead of stimulus-independent private variability.

**Figure 2:**
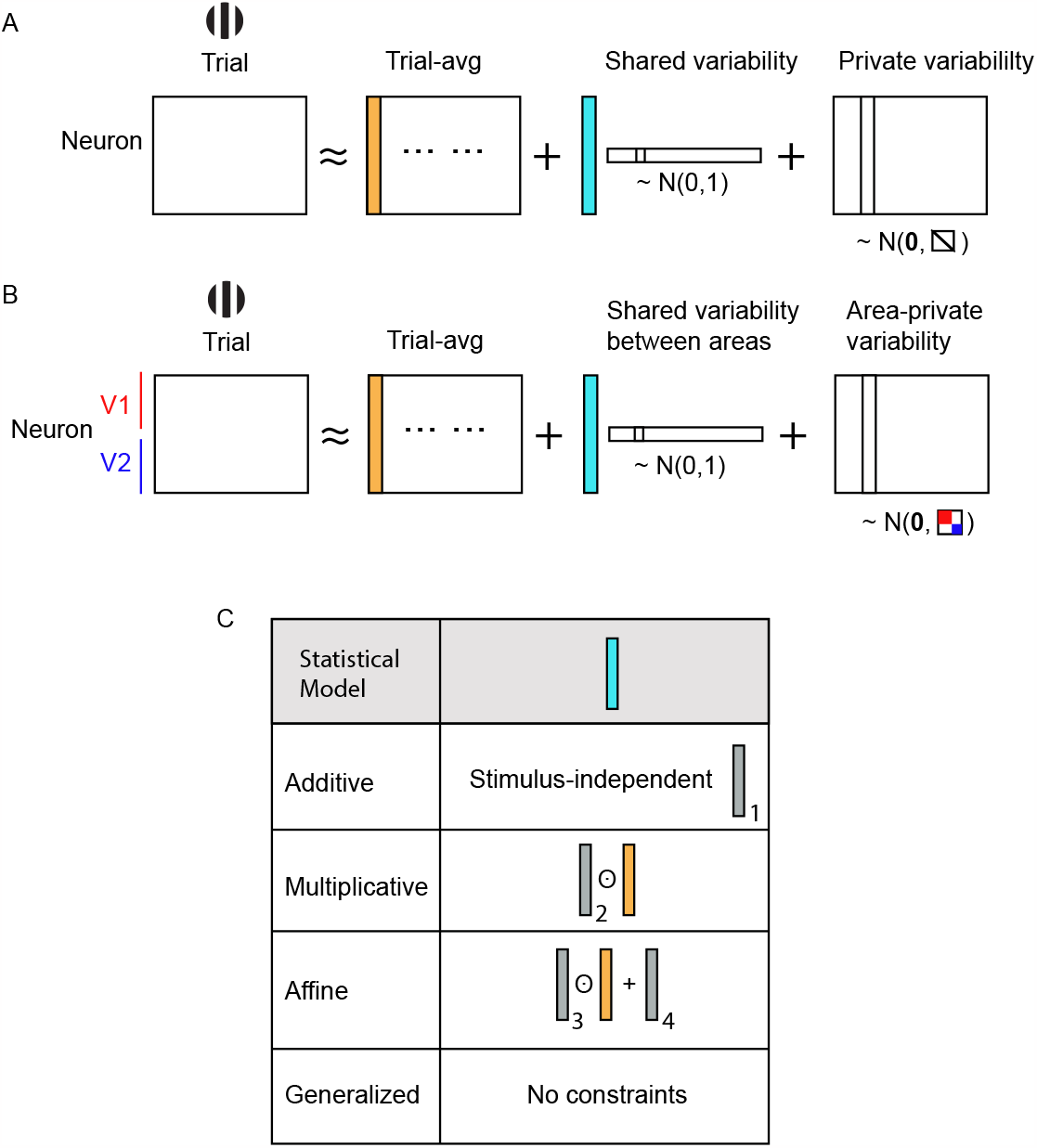
a schematic of the structures of statistical models. A. The statistical model for capturing trial-to-trial variability shared within each brain area. For a given stimulus, we model the single-trial population neuronal responses within a brain area as a sum of 3 terms: the trial-averaged activity (orange column, representing pattern of trial-averaged activity across neurons; repeated identically for each trial), a 1-dimensional Gaussian variable shared across neurons (blue column representing pattern across neurons of shared variability, multiplied by a scalar Gaussian variable for each trial, represented by row; we also studied the case with multi-dimensional shared variability), and private variability that is an independent Gaussian variable for each neuron and trial. B. The statistical model for capturing trial-to-trial variability shared between two brain areas, V1 and V2. Similarly to A, we model the single-trial population neuronal responses as a sum of 3 terms, except that the private variability is not independent across neurons, but is independent between the two brain areas (variability within each area represented by red or blue blocks in illustrated covariance matrix, white indicates values of 0). C. We set up 4 types of statistical models with different constraints on the stimulus-dependence of the shared variability amplitudes (*i .e*., of the blue column). For additive models, the shared variability amplitude is constant across stimuli (but varies across cells, indicated by gray vector). For multiplicative models, the shared variability amplitude is the element-wise product (⊙) of a stimulus-independent vector and the stimulus-dependent trial-averaged population vector. For affine models, the shared variability is the sum of multiplicative and additive components.

We fit the four models to V1 and V2 neuronal responses separately, by maximum likelihood estimation with 5-fold cross validation. We evaluated the fitting performance of each model by two cross-validated measures: 1) the log likelihood of the experimental data given the model (Fig. 3A) 2) the *R*^2^ of the model noise covariance in accounting for the experimental noise covariance (see Methods, Fig. 3B). The log likelihood quantifies overall how well the variability (both shared and private) is captured by the statistical models, whereas the *R*^2^ evaluates specifically how well the shared variability is captured by the statistical models.

**Figure 3:**
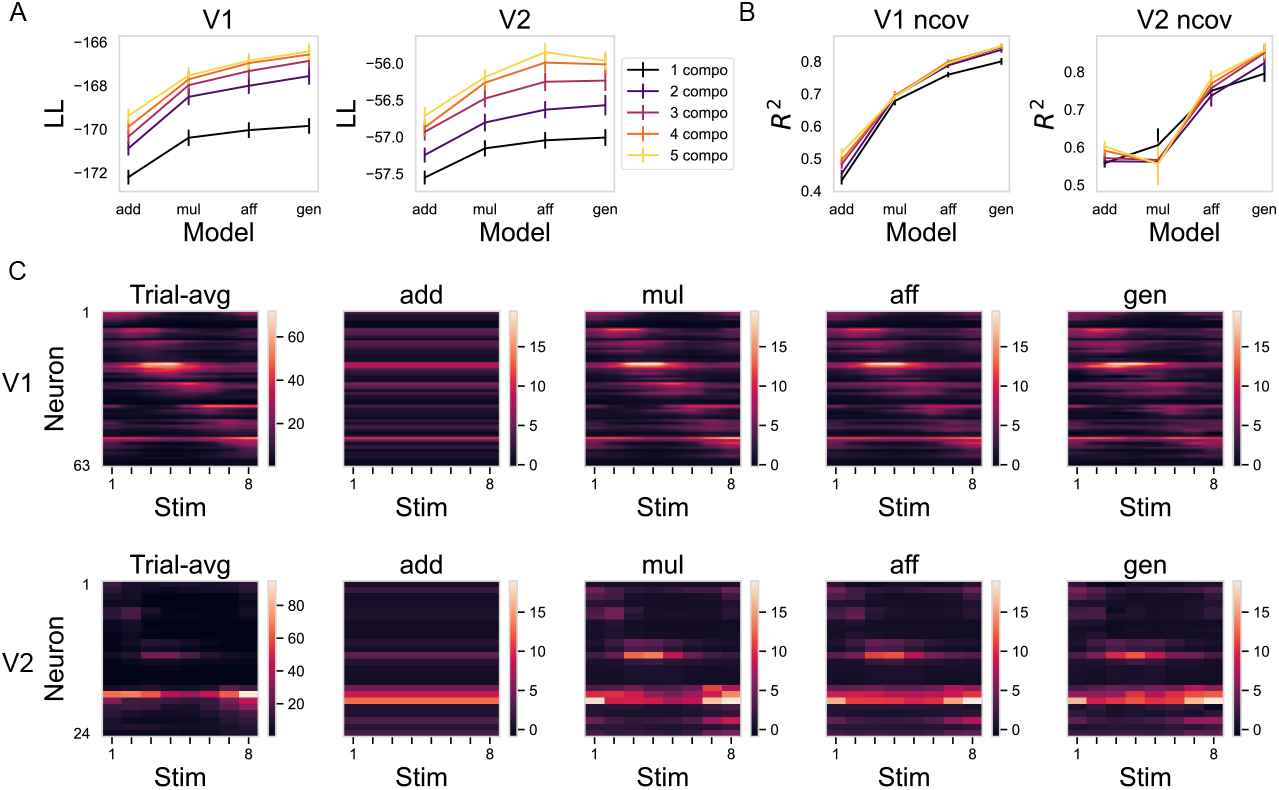
Fitting results of statistical models of experimental session 1. A. The log-likelihood of the test set. Error bars show the standard error of 5-fold cross-validation. Color denotes the number of components of the shared variability. Note that we ordered the statistical models based on their complexity in ascending order from left to right. B. The *R*^2^ of the noise covariance (“ncov”) across stimuli of the test set. Error bars show the standard error of 5-fold cross-validation. Color denotes the number of components of the shared variability. C. Fitted *ϕ* of the 4 statistical models with 1 component in the shared variability are shown against the trial-averaged responses (units Hz) across 8 stimulus orientations. Neurons are ordered according to their preferred orientations. The top row shows the result for V1 neurons. The bottom row shows the result for simultaneously recorded V2 neurons.

All 5 experimental sessions showed the following results consistently (see Fig. 3 for results from one session, and Supplementary Fig. S5 for results from other sessions). First, quantitatively, the generalized model performs the best among all the statistical models, especially in terms of *R*^2^ (Fig. 3A, B). Second, qualitatively, the affine model performs almost as well as the generalized model in terms of capturing how shared variability amplitude changes across orientations (Fig. 3C, inferred *ϕ* exhibits similar patterns for the two models). Third, with less than 5 components in the shared variability, we captured most of the explainable variance in the neural data (Fig. 3, Supplementary Fig. S5). Additionally, inferred private variability has an amplitude proportional to the trial-averaged responses, with proportionality constant *>* 1 (more variable than Poisson noise; Supplementary Fig. S4).

Interestingly, we observed that experimental sessions with high firing rates tend to be better fit by additive models than multiplicative models (Supplementary Fig. S5), suggesting that they exhibit weaker stimulus-dependence of the shared variability. This observation aligns with the theoretical predictions (see box 1) and is supported by the circuit model simulations (Supplementary Fig. S5).

### Generalized affine models explain how shared variability depends on stimulus contrasts and orientations jointly

We predict that affine models will fail to explain how shared variability is modulated by contrasts and orientations jointly. This is because the circuit mechanism (Fig.1B, D), as well as previous experimental and theoretical evidence (M. M. Churchland et al., 2010; Hennequin et al., 2018), suggest that the amplitude of shared variability should decrease with increasing stimulus contrasts, which can’t be captured by affine models. To test this prediction, we recorded from V2 neurons using neuropixel probes while presenting drifting gratings with 8 orientations and 3 contrast levels (see descriptions of dataset 2 in Methods and Supplementary Fig. S2).

Motivated by results of the circuit model simulation (Fig. 1B, D) and fits to experiments varying orientation only (Fig. 3), for a given contrast level, we expect the dependence of shared variability on orientations to be well-approximated by an affine model. Therefore, we proposed a generalized affine model. This model incorporates contrast-specific affine coefficients *α* and *β*, where *ϕ*_*c,r,s*_ = *α*_*c,r*,contrast_*d*_*c,s*_+ *β*_*c,r*,contrast_. We introduced this model because we didn’t have enough data to fit the generalized model. Due to session duration limits and an increasing number of stimuli, we had to reduce the number of trials for each stimulus. Consequently, we only have 100 trials per stimulus. Such a limitation raises concerns about overfitting of complex models. As seen in our results (Fig. 4A, B), even though the generalized models fit the data best in the training set, they perform poorly in the test set.

**Figure 4:**
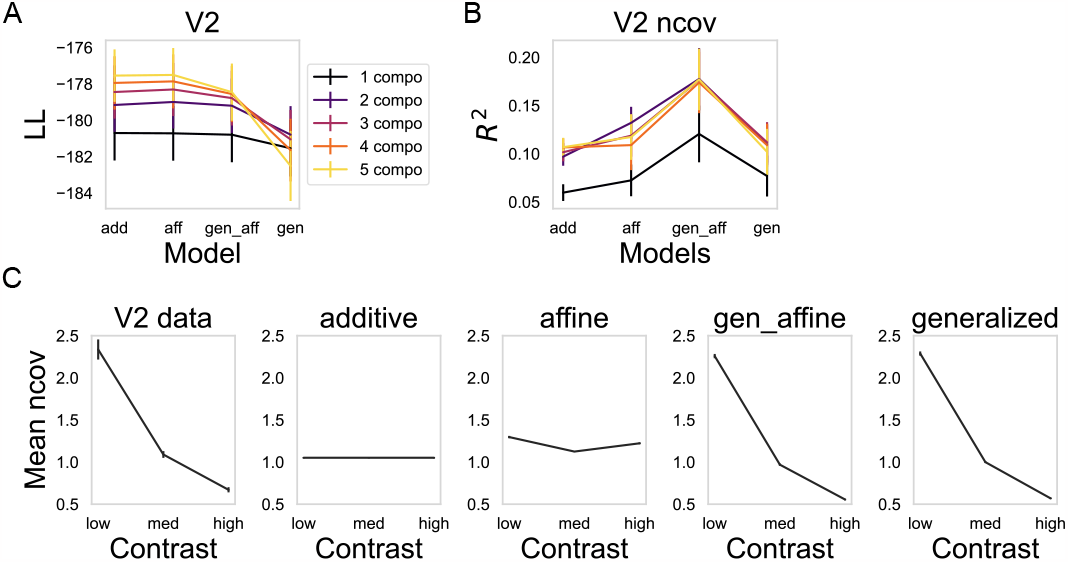
Fitting performances of statistical models on the simultaneously recorded V2 data under stimulation by drifting gratings with varying orientation and contrast. A. The log-likelihood of 4 different statistical models on the test set. Conventions as in Fig. 3A. B. The *R*^2^ of the noise covariance across stimuli of the test set. Conventions as in Fig. 3B. C. The noise covariance averaged over neuron pairs and stimulus orientations is plotted against stimulus contrast levels. The first column shows the noise covariance calculated from V2 data in the test set. Error bars show the standard error of 5-fold cross-validation. The second, third and fourth column show noise covariance calculated from different fitted statistical models with 2 components (“gen aff”: generalized affine).

Judging by the log-likelihood and *R*^2^ of the noise covariance (Fig. 4A, B), the generalized affine model performs the best when both contrasts and orientations are varied. To show that an unmodified affine model indeed fails to capture the modulation of shared variability in this case, we visualized the contrast dependence of the noise covariance, averaged over neuron pairs and stimulus orientations. As predicted by the circuit model, the amplitude of shared variability and the averaged noise covariance decreases with increasing contrast, which is captured by the generalized affine model and the generalized model, and importantly, not by the affine model (Fig. 4C, Supplementary Fig. S7).

### Circuit models predict affine-like variability shared between V1 and V2

So far, our analysis has focused on shared variability within each visual areas. However, a significantly underexplored topic in the existing literature is the structure of variability shared between two inter-connected visual areas. To gain some expectation on how variability shared between V1 and V2 should depend on the stimulus, we simulated a circuit model with two connected ring structures (Fig. 5A), where each ring represents one visual area. We assumed that connectivity from area 1 to area 2 is similar to connectivity within each area, except without inhibitory projections (Fig. 5A).

**Figure 5:**
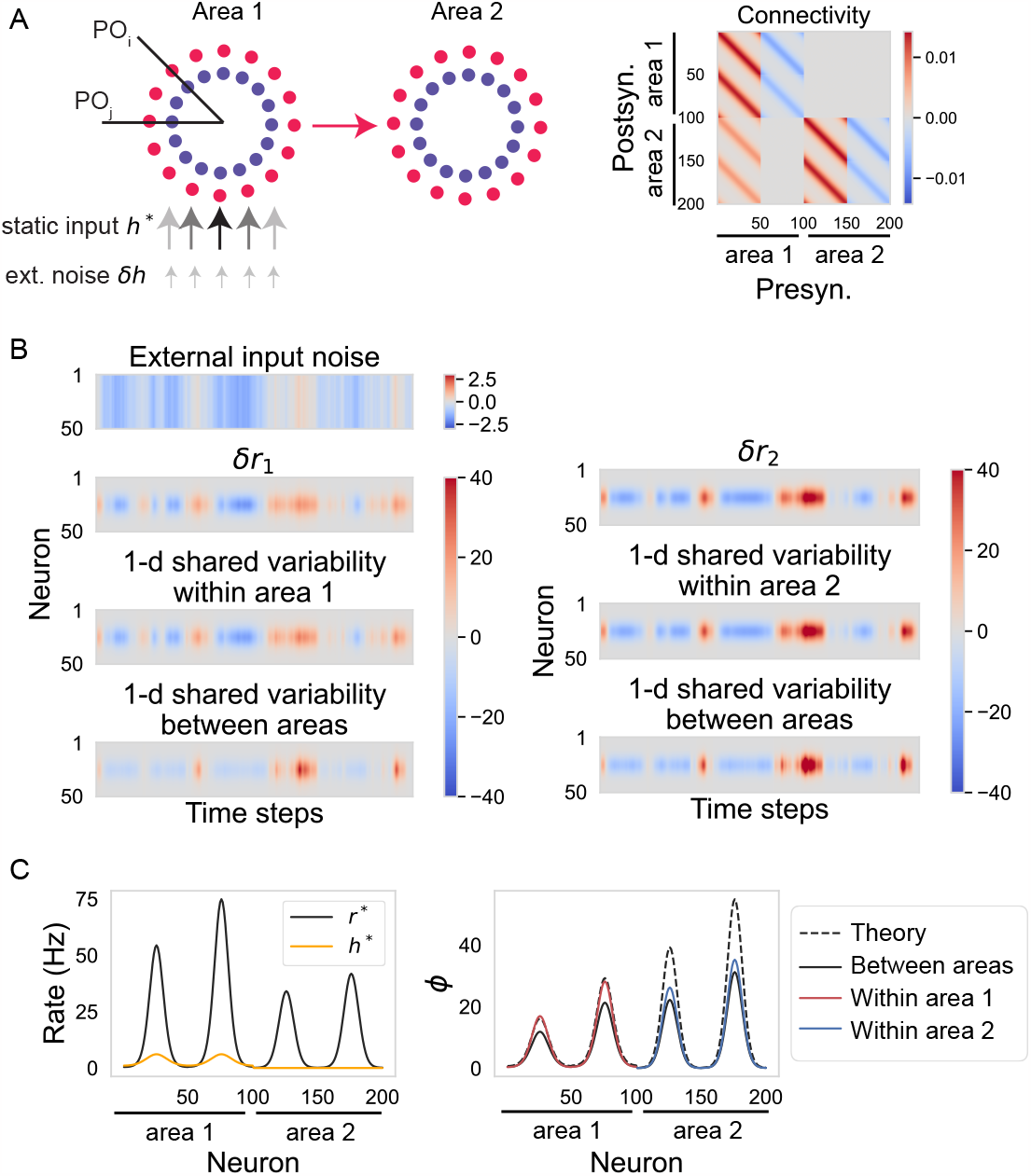
Connected ring models predict affine shared variability between brain areas. A. Left: Schematic of two-ring models, with each ring representing one brain area. Between-area connections are excitatory and unidirectional, from area 1 to area 2. Area 1 receives an excitatory external input. Right: the connectivity strengths between the model neurons. The first 100 neurons are 50 excitatory and 50 inhibitory neurons in area 1, the last 100 neurons are in area 2. B. Residual activity of excitatory neurons in area 1 (left) and area 2 (right). Left: (from top to bottom panels) external input noise *δh* received by the excitatory neurons in area 1; residual activity *δr* of the excitatory neurons in area 1; the dominant 1-dimensional variability shared within area 1; the dominant 1-dimensional variability shared between two areas. Right: similar to figures on the left but for area 2. Note that area 2 does not receive any external input noise directly. C. Left: the external static input *h*^*^ and the time-averaged rate responses *r*^*^ of all the simulated neurons in the connected ring model. Right: the 1-dimensional shared variability amplitudes of all the simulated neurons. Dashed: analytical prediction from eq.5. Red: amplitude of variability shared within area 1. Blue: amplitude of variability shared within area 2. Black: amplitude of variability shared between two areas. The gap between analytical prediction and numerical simulation in area 2 is due to a violation of the small input noise assumption.

We simulated the response of neurons in the connected ring network to changing stimulus orientation (Fig. 5B). Area 1 received external inputs that consist of an orientation-tuned static external input and 1-dimensional external input noise. To calculate the amplitude of 1-dimensional variability shared within each area, we looked at the dominant component found by applying singular value decomposition (SVD) to the residual activity of neurons within each ring, which is equivalent to the generalized model in the absence of private noise. To calculate the amplitude of 1-dimensional variability shared between areas, we looked at the dominant component found by applying probabilistic canonical correlation analysis (pCCA) (Bach and Jordan, 2005) to the set of residual responses of the two areas. This is equivalent to a generalized joint model.

The variability shared between areas 1 and 2 consists of a pattern within each area (Fig. 2B). Within area 1, this shared variability may form a similar pattern to the purely within-area shared variability (Fig. 5) or a distinct pattern (Supplementary Fig. S8), depending on the chosen connectivity between area 1 and area 2 (because area 2 does not receive any external noise, its within-area and between-area noise always resemble one another). Essentially, if the connectivity from area 1 to area 2 has a reasonably strong component along the patterns in *δr*_1_, these patterns in *δr*_1_ will propagate to area 2, resulting in a between-area shared variability pattern that resembles the within-area-1 shared variability (Fig. 5 B, C). Conversely, if the between-area connectivity and the *δr*_1_ patterns are close to orthogonal, area 1’s pattern of between- and within-area shared variability may differ (see an example simulation in Supplementary Fig. S8). It is reasonable to assume that the trial-averaged activity in V1 has a strong component along the feedforward connectivity; otherwise, V2 wouldn’t be strongly activated. Building upon our previous analysis, which indicates that the dominant pattern of shared variability within V1 aligns closely with trial-averaged activity and is affine (Fig. 3C), we predict that shared variability between visual areas will exhibit a similar orientation-dependent pattern to shared variability within each visual area and can be effectively captured by affine models.

### Affine joint models parsimoniously explain variability shared between V1 and V2 across stimulus orientations

We now characterize the stimulus-dependence of variability shared between V1 and V2, in responses to stimuli of varying orientation at a fixed (100%) contrast. We use four statistical models like the four used previously, modified for the two-area case (Fig.2B).

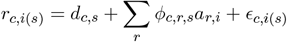

The key differences are the following: first, we fit the statistical models to V1 and V2 data jointly as opposed to separately, thus, we refer to these statistical models as “joint statistical models”; second, we assume the private variability *ϵ*_*c,i*(*s*)_ is private to each brain area instead of private to each neuron (and thus can be shared within an area). Specifically, we assume that on each trial *i* for a given stimulus *s, ϵ*_*i*(*s*)_, which is a vector with length of the total number of neurons in V1 and V2, is a sample taken from a multivariate Gaussian distribution 𝒩 (0, Σ_*s*_). We assume that Σ_*s*_ is block diagonal, having non-zero entries only for within-V1 or within-V2 covariances. Here, *ϕ*_*c,r,s*_ denotes the amplitude of the *r*^*th*^ component of shared variability between areas for neuron *c*, stimulus *s*.

As before, we fit joint statistical models to V1 and V2 data by maximum likelihood estimation with 5-fold cross-validation, and we evaluated the performances of the joint statistical models with the previously mentioned two measures. Without overfitting, we should expect to see that the generalized model performs the best (see Supplementary Fig. S9 for fitting performances of joint statistical models on surrogate data with known ground truth).

All 5 experimental sessions showed the following results consistently (Fig. 6, Supplementary Fig. S10). Quantitatively, the generalized joint model performs the best in terms of capturing the shared variability between V1 and V2, slightly outperforming the affine joint model in terms of *R*^2^. Qualitatively, the affine joint model inferred a similar stimulus-dependence pattern for the shared variability amplitude as the generalized joint model (Fig. 6C). As in (Semedo, Zandvakili, et al., 2019; Semedo, Jasper, et al., 2022), we found that 1 or 2 components for variability shared between areas are enough for capturing all the explainable variance in the experimental data. We conclude that affine joint models parsimoniously explain how variability shared between V1 and V2 varies across orientations.

**Figure 6:**
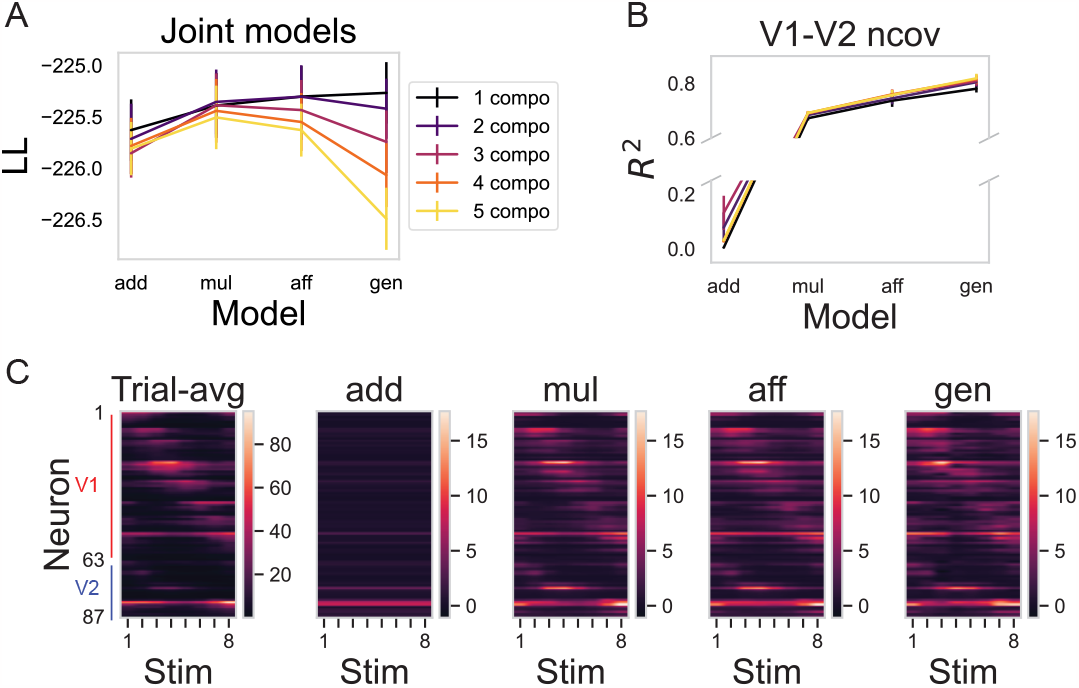
Fitting results of joint statistical models for both V1 and V2 of experimental session 1. A. The log-likelihood of 4 different joint statistical models on the test set. Conventions as in previous figures. B. The *R*^2^ of the noise covariance between V1 and V2 across stimuli of the test set. Conventions as in previous figures. For visualization purposes, we omitted part of the y axis. C. Fitted *ϕ* of the 4 joint statistical models with 1 component are shown against the trial-avg rates (units Hz) across stimuli with 8 orientations. Neurons are ordered according to their preferred orientations within each brain area.

Additionally, we found that the amplitude of variability shared between areas is smaller than that within each area, as expected from the circuit model simulation (Fig. 5C, Supplementary Fig. S11, Supplementary Fig. S12).

## Discussion

We studied the stimulus dependence of shared variability within and between V1 and V2. When only stimulus orientation is varied, affine models effectively describe this stimulus dependence. However, when both orientation and contrast are varied, affine models must be modified to include contrast-specific coefficients to explain the suppression of shared variability with increasing contrast. Study of a recurrent neural circuit with power-law activation function, subjected to correlated stimulus-independent external input noise, suggests a mechanistic rationale for these statistical models.

### Novelty of our work

In the previous literature, statistical models partition neural variability in one of two ways. First, motivated by the fact that spike count variance grows faster than its mean across stimuli, neural variability is partitioned into firing rate variability and spiking variability (usually modeled as Poisson noise). Second, as assumed by our work, neural variability is partitioned into shared and private variability. In general, the first method is used for explaining the variability of individual neurons (A. K. Churchland et al., 2011; Goris, Movshon, and E. P. Simoncelli, 2014; Zhu and Wei, 2023), whereas the second method is used for explaining variability in simultaneously recorded neuronal populations (Rabinowitz et al., 2015; Lin et al., 2015; Arandia-Romero et al., 2016). In our study, we extend the findings in the second framework in four key directions: 1) by comparing three previously proposed forms of modulation (additive, multiplicative, affine) to an unrestricted form (generalized), we provide evidence that affine models offer a parsimonious explanation for how shared variability is modulated by stimulus orientations; 2) we establish a direct link between the statistical models and a neural circuit model, offering a straightforward mechanism for the observed affine shared variability; 3) we identified an alternative form of stimulus-dependence of shared variability (generalized affine), which arises when stimulus strength (contrast) is varied; 4) we broaden the framework to explain variability shared between two connected brain areas, demonstrating that variability shared between V1 and V2 also exhibits an affine pattern across stimulus orientations.

### Implications for stimulus coding

We find that the dominant component of shared variability within or between V1 and V2 closely aligns with the trial-average activity. Therefore, most of shared variability is orthogonal to the direction in which a slight change in stimulus orientation modulates population activity but aligns with the direction in which a slight change in the contrast level modulates population activity. This suggests that shared variability would have limited impact on the coding of orientation but could be detrimental to the coding of contrast (Moreno-Bote et al., 2014). However, as the contrast level increases, the influence of shared variability on coding diminishes due to its decreasing amplitude.

### Implications for circuit structure

Investigations of circuit mechanisms give insight into how external inputs can modulate neural variability. As shown in box 1 and discussed in Doiron et al., 2016, the alteration of patterns of trial-averaged activity induced by changes in stimuli will alter the patterns of variability, even when external input noise remains stimulus-independent. Since behavioral variables and task conditions also alter trial-averaged activity levels, it is not surprising to observe that they too induce modulations in neural variability.

The circuit model also shows that, when firing rates become sufficiently high, and if external input noise is stimulus-independent, then neural variability should become stimulus-independent (*δr*(*t*) ≈ *W*^−1^*δh*(*t*), see box 1) and therefore well fit by an additive model. Indeed, we have found that in some sessions with particularly high firing rates, an additive model gives the best fit (Supplementary Fig. S6). In the other extreme, when the firing rates are sufficiently low to render effective connectivity very weak, the variability becomes independent of the recurrent connectivity, being determined by effective gains and external input noise (*δr*(*t*) ≈ *Fδh*(*t*), see Box 1).

In most scenarios, recurrent connectivity affects the pattern of neural variability through eq. 5, which suggest that the variability is not necessarily always well-described by affine models. Our simulation demonstrated how affine models can fall short, particularly when we vary the contrast level of the stimulus, which is supported by experimental data analysis (Fig. 4). Furthermore, the circuit model predicts that the amplitude of shared variability can exhibit an “M”-shaped pattern across orientations in response to sufficiently strong tuned input (Fig. 1), as has been observed in area MT (Ponce-Alvarez et al., 2013). However, our datasets are limited to only 8 orientations spanning a full 180-degree range (see Methods). Consequently, our statistical models may not discern subtle variations, such as distinguishing between an “M”-shaped and a bump-shaped modulation. Alternatively, it is possible that even at the highest contrast level, V1 may not receive sufficiently strong tuned input to elicit the “M” shape. Future experiments with finer resolution within the stimulus space could resolve this issue.

In summary, our investigations of circuit mechanisms shed light on how external input can modulate neural variability. Statistical models effective in V1 and V2 may not be optimal for other brain areas, due to variations in recurrent connectivity. Nonetheless, insights into the two extreme cases of very low or very high firing rates are likely to be more general.

### Implication for communication between brain areas

Previous work has shown that the communication subspace between V1 and V2, when stimulated by oriented drifting gratings, is limited to just 1 or 2 dimensions out of perhaps 5 dimensions of variability within each area (Semedo, Zandvakili, et al., 2019; Semedo, Jasper, et al., 2022). Such low dimensionality is unexpected for early processing stages like V1 and V2 in the visual hierarchy. However, the shared variability between V1 and V2 varies across stimuli, as shown by our work and a previous study (Semedo, Zandvakili, et al., 2019), so the combined communication subspace under different stimuli can be higher dimensional than only considering a single stimulus.

## Acknowledgments

We thank Jeff Johnston, Salomon Zev Muller, Ashok Litwin-Kumar and members of our group for comments and discussions. We thank Amin Zandvakili for recording dataset 1 and Aravind Krishna for recording dataset 2. This work was supported by Simons Foundation 543071, Simons Foundation 542999, NSF 1707398, and Gatsby Charitable Foundation GAT3708.

## Methods

### Neural recording and visual stimulation

Animal procedures have been reported in previous work (Smith and Kohn, 2008; Zandvakili and Kohn, 2015). Briefly, animals (macaca fascicularis, 2-5 years old) were anesthetized with ketamine (10 *mg/kg*) and maintained on isoflurane during surgery. All recordings were performed under sufentanil (6-18 *μg/kg/hr*) anesthesia. Vecuronium bromide (150 *μg/kg/hr*) was used to prevent eye movements. All procedures were approved by the IACUC of the Albert Einstein College of Medicine.

Dataset 1 has been reported previously (Zandvakili and Kohn, 2015; Semedo, Zandvakili, et al., 2019) and is publicly available under https://crcns.org/data-sets/vc/v1v2-1/. Recordings in V1 were performed using a Utah Array (96 channels, Blackrock Neurotech, USA) and recordings in V2 were performed using a set of tetrodes (Thomas Recording, Germany). Stimuli (full contrast drifting gratings, 8 orientations in steps of 22.5 deg, 1 *cyc/d*, drift rate of 3-6.25 Hz; 2.6-4.9 deg in diameter) were presented on a gamma corrected CRT screen (1024-796 pixel, 100 Hz refresh). Each stimulus was presented 400 times for 1.28 s preceded by an interstimulus interval of 1.5 s. Voltage snippets that exceeded a user defined threshold were digitized and sorted offline. The recorded V1-V2 populations had overlapping receptive fields.

Dataset 2 was recorded using four Neuropixel 1.0 (IMEC, Belgium) probes inserted in a rhomboid arrangement (see Supplementary Fig 1) with the most anterior sites being roughly 3 mm posterior of the lunate sulcus. Data was recorded with a ‘linear configuration’ resulting in 384 sites distributed over 7.6 mm of the probe shaft. Area boundaries for V1 and V2 were determined by a combination of change in receptive field position and size and histological reconstruction of the electrode track. Stimuli (drifting gratings, 8 orientations in steps of 22.5 deg, 2 *cyc/d*, drift rate of 4 Hz; 8 deg in diameter; 15%, 50% and 100% Michelson contrast with *L*_*min*_ = 0 *cd/m*^2^ and *L*_*max*_ = 80 *cd/m*^2^). Each stimulus was presented 200 times for 0.5 s preceded by a 0.5 s inter stimulus interval. Voltage traces were digitized using spikeGLX (https://billkarsh.github.io/SpikeGLX/) and sorted offline using Kilosort 2.5 (Steinmetz et al., 2021) and Phy2 (https://github.com/cortex-lab/phy). The V1 and V2 populations had partially overlapping receptive fields.

For dataset 2, we didn’t show the fitting results of statistical models for V1, due to lack of sufficient data.

### Data preprocessing

For dataset 1, for each neuron and trial, we counted the spikes during the stimulus excluding the first 100 ms after stimulus onset. For dataset 2, for each neuron and trial, we counted the spikes during the stimulus. Additionally, we concatenated 2 trials as 1 trial to increase the number of spikes per trial, ensuring that the Gaussian assumption in our statistical models remains valid. We ended up with 100 trials per stimulus.

We quantified how well-tuned each neuron is by calculating the orientation tuning index (OTI). The neuron’s tuning curve is just the trial average of its response at each orientation. OTI is computed as

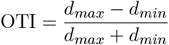

where *d*_*max*_ and *d*_*min*_ are the maximum and minimum of the tuning curve, respectively. When fitting the statistical models, we excluded neurons with 0 spikes over trials for any given stimulus and excluded neurons with a fano factor *>* 4 in dataset 2 to avoid unstable units. Furthermore, we excluded neurons with OTI *<* 0.35 for both datasets. In dataset 1 (5 recording sessions), our study included 359 out of 564 units in V1 and 119 out of 147 units in V2. In dataset 2 (1 recording session), our study included 46 out of 132 units in V1 and 74 out of 191 units in V2.

### Fitting statistical models

We have 5 statistical models for describing neural variability within each brain area: additive, multiplicative, affine, generalized affine and generalized models. We fit the models separately to V1 and V2 data with 5-fold cross-validation. At each fold, we split 400 trials into a training set of 320 trials and a test set of 80 trials, chosen at random. We fit the model to the training set with maximum likelihood estimation, and evaluated the fitting performance by the log likelihood of the test set.

As noted in the Results, the generalized model is equivalent to factor analysis applied separately to the residual responses to each stimulus. To fit generalized models, we used the python class “sklearn.decom position.FactorAnalysis” with the default initialization parameters.

For the other 4 models, careful initialization was required for a fair comparison between models, because the log likelihood of our statistical models can have many local maxima. To initialize the parameters in the other 4 statistical models, we fit a naive version of the additive model with stimulus-independent private variability by applying factor analysis to population residual responses to all stimuli. Then the models were initialized as follows:

Initialization of the additive model:

- Initialize *ϕ*_*c,r*_ with the fitted *ϕ*_*c,r*_ of the naive additive model.
- Initialize 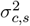 with the fitted 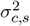 of the generalized model.

Initialization of the affine model:

- Initialize *α*_*c,r*_ as all zeros.
- Initialize *β*_*c,r*_ as the fitted *ϕ*_*c,r*_ from the naive additive model.
- Initialize 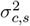 with the fitted 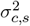 of the generalized model.

Initialization of the multiplicative model:

- Initialize *α*_*c,r*_ as fitted *α*_*c,r*_ from the affine model.
- Initialize 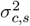 with the fitted 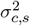 of the generalized model.

Initialization of the generalized affine model:

- Initialize *α*_*c,r*,contrast_ as all zeros.
- Initialize *β*_*c,r*,contrast_ across all contrast levels as the fitted *ϕ*_*c,r*_ from the naive additive model.
- Initialize 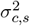 with the fitted 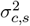 of the generalized model.

For additive, multiplicative, affine, and generalized affine models, we optimized the parameters by gradient descent with the Adam optimizer. We chose the learning rate to be 5e-3 or 1e-2 with a 0.96 decay rate. We stopped the optimization after the log likelihood of the training set converges or at the 20, 000^th^ iteration.

### Fitting joint statistical models

We have 4 joint statistical models for describing neural variability between two brain areas: additive, multiplicative, affine and generalized joint models (we did not use generalized affine because contrast was not varied in these studies). We fit the models jointly to V1 and V2 data with 5-fold cross-validation.

We wanted to characterize the shared variability between V1 and V2, separate from shared variability within each area. This means that, in fitting a joint model to V1 and V2, the private variability is private to each area (can be correlated within each area), rather than being private to each neuron. For this reason, we cannot use the methods we applied to each single area. Instead, to fit generalized joint models, we applied probabilistic canonical correlation analysis (pCCA) to the population residual responses from V1 and V2 to each stimulus separately (as in Semedo, Jasper, et al., 2022). Similarly to the fitting process of within-area statistical models, we initialized the parameters in the additive, multiplicative and affine joint models with the fitted parameters from the generalized joint models or naive additive joint models with stimulus-independent area-private variability. Naive additive joint models were fitted by applying pCCA to the population residual response from V1 and V2 to all stimuli.

### Evaluating model performance

We quantified how well the statistical models explain all the variance in the data by cross-validated log likelihood, as previously described (see Methods section: fitting statistical models). We also quantified how well the statistical models explain the shared variability in the data by evaluating the cross-validated *R*^2^ of the noise covariance matrix across all stimuli. Specifically, we computed the noise covariance matrix from the test set for each stimulus and concatenated the upper triangular elements of these matrices across all stimuli. We then computed the *R*^2^ describing how well this concatenated vector was predicted by the statistical models fit to the training set. The noise covariance between neuron *c*_1_ and neuron *c*_2_ of a given stimulus s from the test set is calculated using the formula: 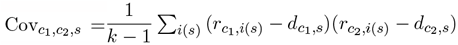, where *k* is the number of trials per stimulus in the test set, *i*(*s*) is the trial index during stimulus *s, r* is the single-trial response, *d* is the trial-averaged response. The noise covariance of a given stimulus *s* from the fitted statistical model is calculated as 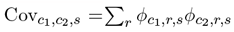. Note that we chose to look at noise covariance instead of noise correlation here, because noise correlation is also influenced by the stimulus-dependent private variability, not just the shared variability.

Similarly, we quantified how well the joint statistical models capture variability shared between V1 and V2 by cross-validated *R*^2^ of the noise covariance between areas. Here, noise covariance is defined for all possible neuron pairs, where each neuron pair consists of one V1 neuron and one V2 neuron. The noise covariance is calculated using the formula described earlier, except that *c*_1_ is the index of neurons in V1, and *c*_2_ is the index of neurons in V2. We evaluated the *R*^2^ for all elements in the noise covariance matrix between V1 and V2 across all stimuli.

### Neural circuit model

We simulated a rate-based network with a ring architecture. The network contained 50 excitatory and 50 inhibitory units. The circuit dynamics was governed by:

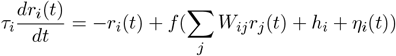

*r*_*i*_ denotes the rate for neuron *i, τ*_*i*_ is the time constant for neuron *i, W*_*ij*_ is the synaptic strength from neuron *j* to neuron *i, h*_*i*_ is the external static input received by neuron *i, η*_*i*_ is the external input noise received by neuron *i*. The activation function *f* is defined as: 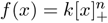.

The connectivity matrix *W* is set to be translational invariant, and the strength of connections between two neurons follows a circular Gaussian of the difference in their preferred orientations:

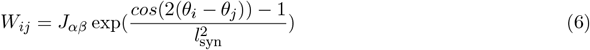

where *α, β* ∈ {*E, I*}, *J*_*αβ*_ is a scaling constant for synaptic strength between E and I neurons, set so that the sum of incoming E and I weights onto each E and I neuron matched the values of *W*_*αβ*_. *θ*_*i*_ denotes the preferred orientation of neuron *i. l*_*syn*_ sets the length scale over which synaptic weights decay.

The external static input *h*_*i*_ representing stimulus-related input. It is constant and follows a circular Gaussian function with an added baseline:

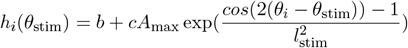

*θ*_stim_ denotes the orientation of the visual stimulus. The neuron with preferred orientation matching the stimulus orientation receives the strongest external static input, and *l*_stim_ sets the length scale over which the external static input decay. *b* denotes the constant baseline. *c* denotes the contrast level. *A*_max_ denotes the maximum amplitude of external static input.

The external input noise *η*_*i*_(*t*) is modeled as a multivariate Ornstein-Uhlenbeck process: 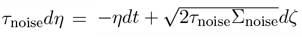 Here we chose the external input noise to be perfectly correlated across neurons so that Σ_noise_ is a matrix with all the elements equals to a chosen variance 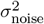. In other words, we had 1-dimensional external input noise that is identical across neurons.

To numerically calculate the amplitude of shared variability *ϕ*, we applied SVD to the population residual responses. This is equivalent to factor analysis given that there is no private variability in the rate model. *ϕ* is calculated as the standard deviation of the dominant SVD component.

To study variability shared between two connected brain areas, we simulated two ring circuits with a feedforward connection from one to the other (Fig. 5). To numerically calculate the amplitude of variability shared between the ring circuits, we applied pCCA to the population residual responses from the two ring circuits. Here, *ϕ* is calculated as the standard deviation of the dominant pCCA component.

**Table 1:** Parameters used in the circuit model in Fig. 1.

|  |  |  |
| --- | --- | --- |
| $N_E$ | 50 | - |
| $N_I$ | 50 | - |
| $\tau_E$ | 20 | ms |
| $\tau_I$ | 10 | ms |
| $k$ | 0.3 | $mV^{-n} \cdot s^{-1}$ |
| $n$ | 2.5 | - |
| $W_{EE}$ | 0.25 | $mV \cdot s$ |
| $W_{IE}$ | 0.24 | $mV \cdot s$ |
| $W_{EI}$ | 0.13 | $mV \cdot s$ |
| $W_{II}$ | 0.1 | $mV \cdot s$ |
| $\tau_{noise}$ | 500 | ms |
| $\sigma_{noise}^2$ | 0.5 | $mV^2$ |
| $\ell_{syn}$ | 45 | deg. |
| $\ell_{stim}$ | 60 | deg. |
| $b$ | 0.1 | mV |
| $A_{max}$ | 20 | mV |

## Supplementary Information

**Figure S1:**
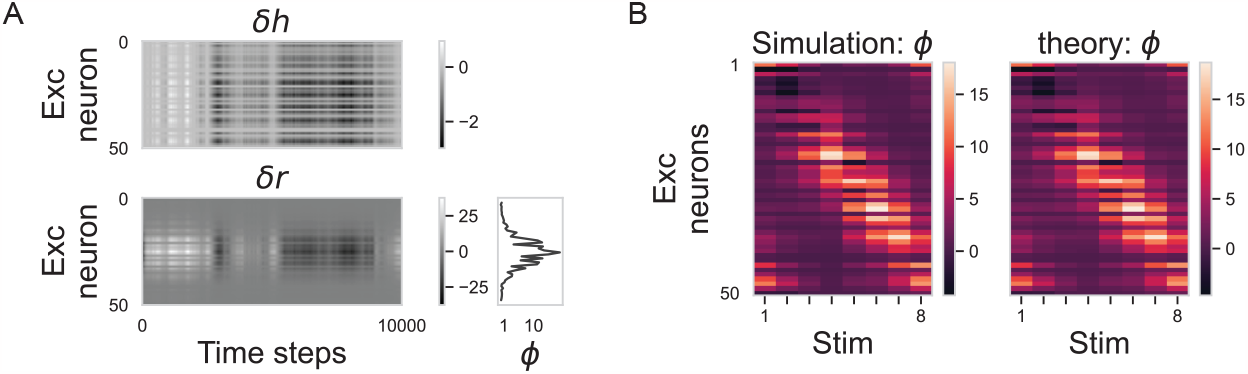
Supplementary to Fig.1, we present model results with heterogeneous external noise. In this case, how the amplitude of shared variability *ϕ* change across orientations is not the same as how it changes across neurons. Therefore, we need to directly visualize how *ϕ* change across neurons and orientations. A. (Top) perfectly correlated external noise with heterogeneous amplitude among recipient neurons and (bottom) the corresponding residual responses. On the right, we plotted *ϕ* across neurons. B. *ϕ* for excitatory model neurons across stimuli with 8 different orientations, calculated from simulations and theoretical predictions (see eq.5).

**Figure S2:**
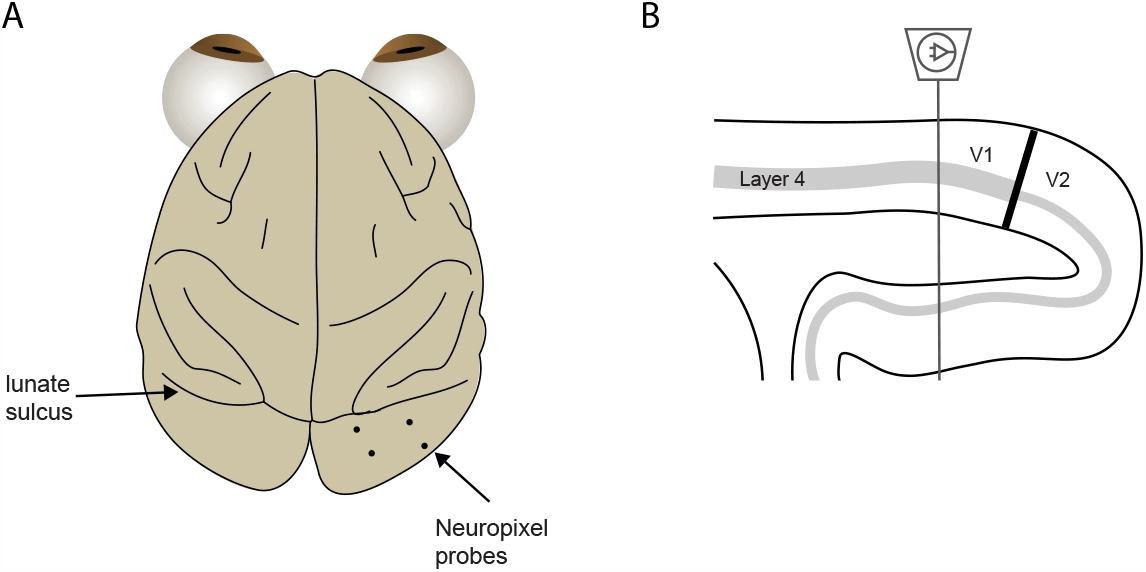
Experimental setup for the neuropixel dataset. A. Arrangement of the four Neuropixel probes on the cortical surface. B. The schematic shows a sagittal section of occipital cortex and one Neuropixel probe.

**Figure S3:**
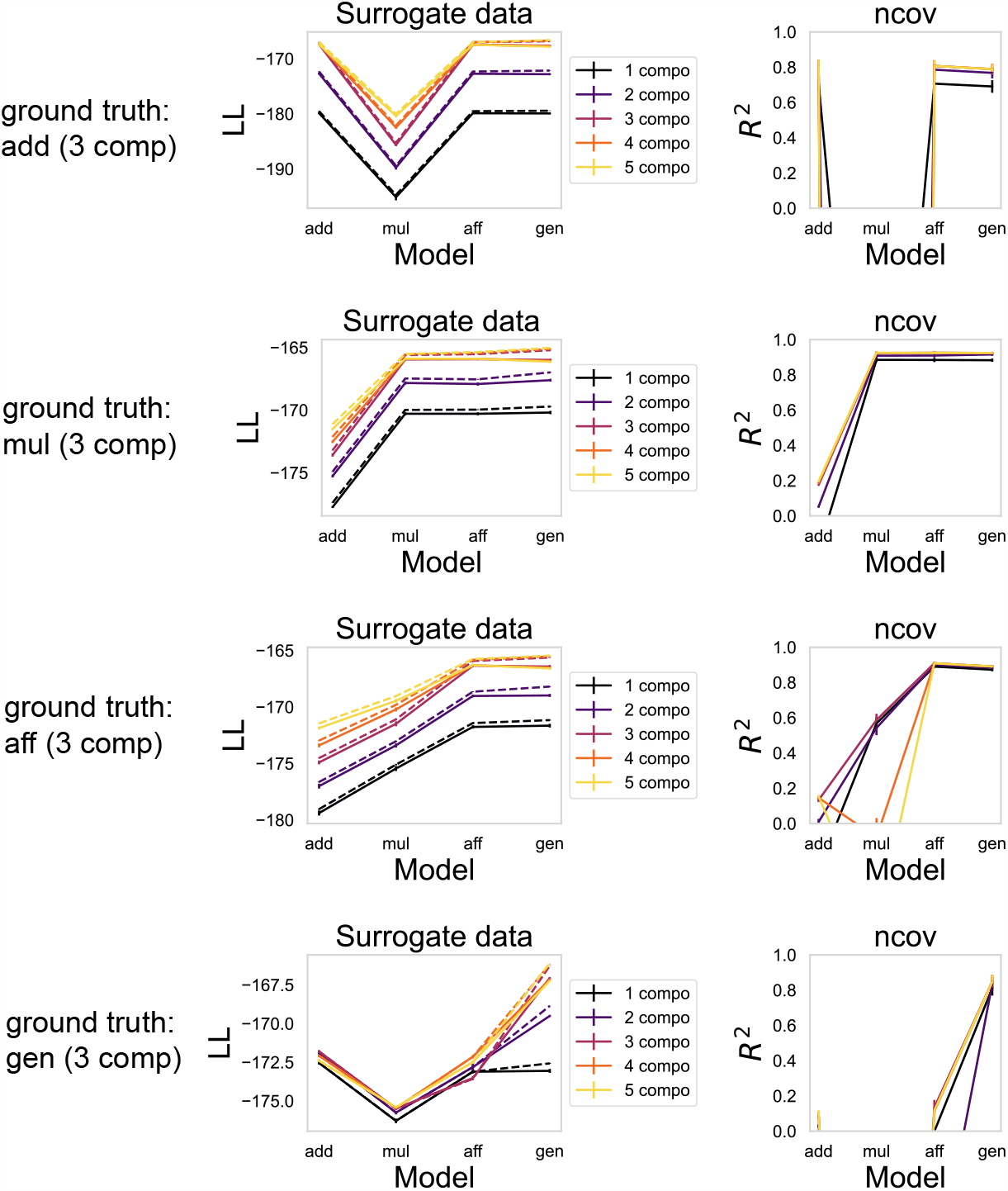
Fitting performances of statistical models on surrogate data with known ground truth. We generated surrogate data (50 neurons × 400 trials × 8 orientations) with similar statistics to the experimental data we have: 1) each neuron has a bell-shaped orientation tuning curve, with trial averaged response span from 20 to 100 Hz across orientations 2) on each trial, shared variability is simulated as a sample taken from a 3-dimensional standard Gaussian distribution. For a given neuron and stimulus orientation, when the ground truth is “add”/”mul”/”aff”, the amplitude of shared variability follows an additive/multiplicative/affine function of trial-averaged responses; when the ground truth is “gen”, the amplitude of shared variability follows a periodic function with 3 peaks across orientations. 3) On each trial, for a given neuron and stimulus orientation, private variability is simulated as a sample taken from a Gaussian distribution with zero-mean and a variance equal to its trial-averaged response.

Similar as in Fig. 3, fitting performances of statistical models were evaluated in two ways: cross-validated log likelihood (left column) and cross-validated *R*^2^ of noise covariance (right column). Dashed line shows log likelihood of the training set, solid line shows log likelihood of the test set. Statistical models with the 3 components and the correct assumption for the shared variability achieved the best performance with the least number of parameters.

**Figure S4:**
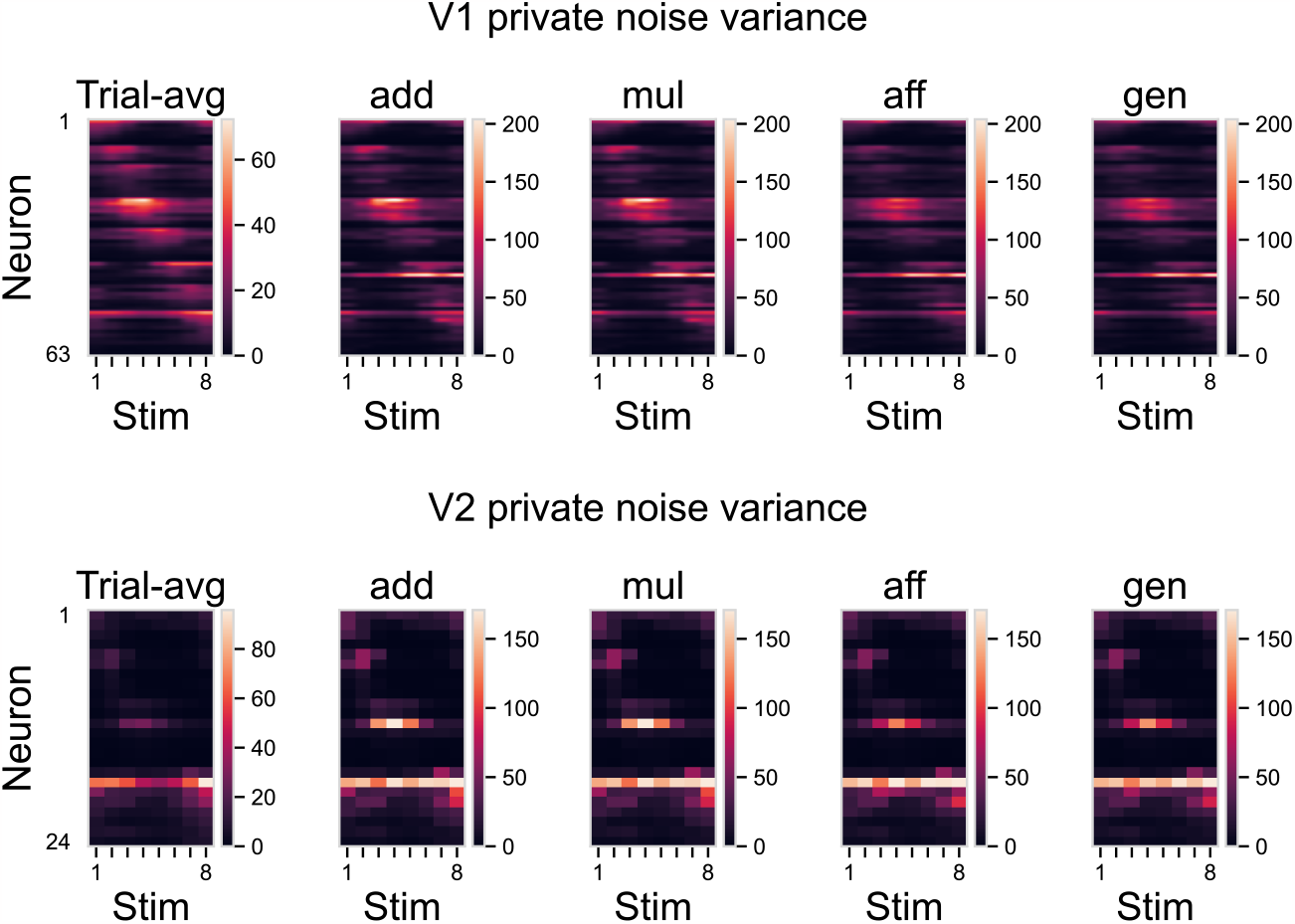
The variance of private variability in the fitted statistical models to session 1 with 1 component for shared variability. The inferred private variability has a variance proportional to the trial-averaged responses.

**Figure S5:**
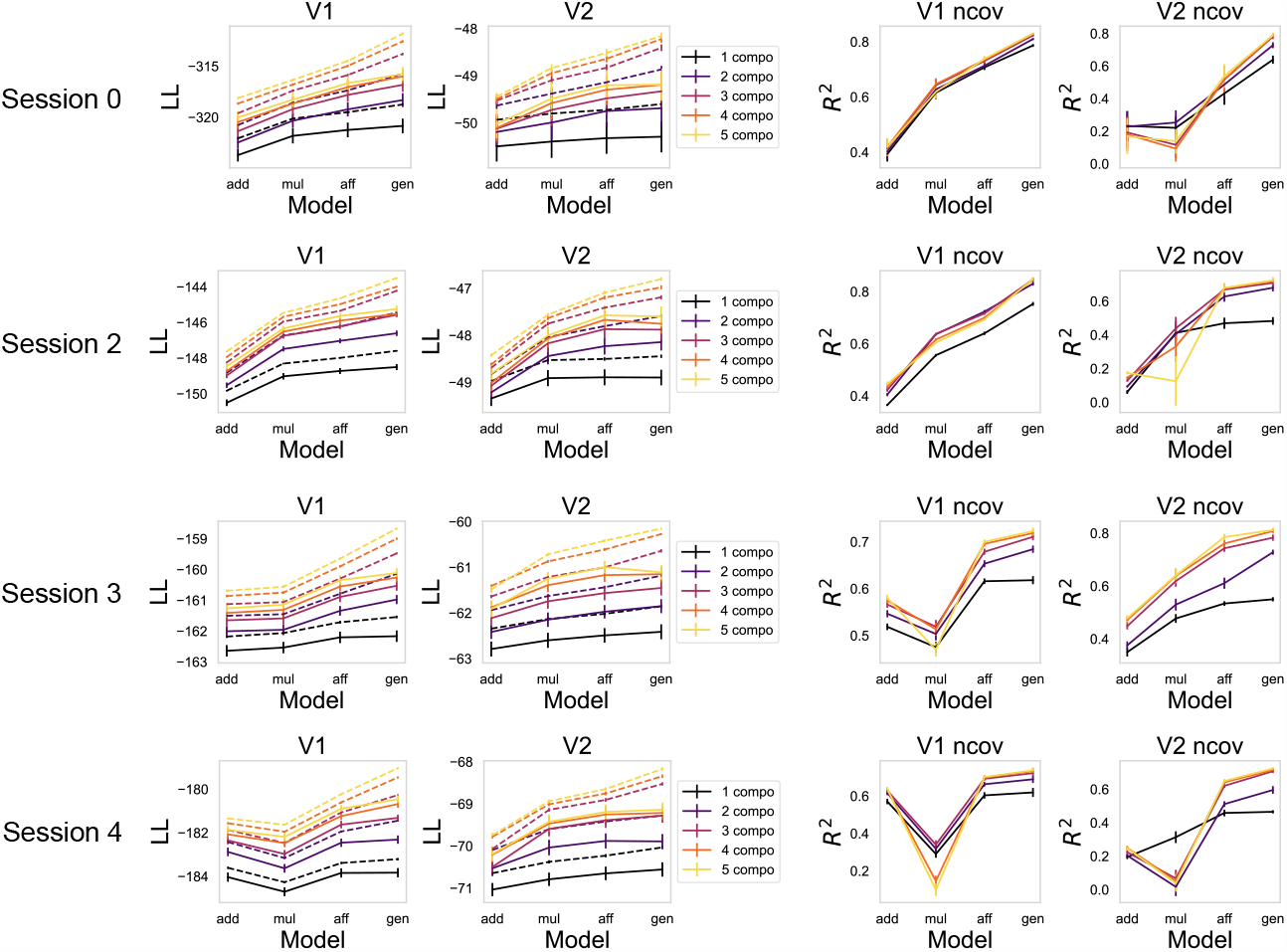
Fitting performances of statistical models on V1 and V2 data for session 0,2,3,4 in the dataset 1. Similar as in Fig. 3, fitting performances of statistical models were evaluated in two ways: cross-validated log likelihood and cross-validated *R*^2^ of noise covariance. Dashed line shows log likelihood of the training set, solid line shows log likelihood of the test set.

**Figure S6:**
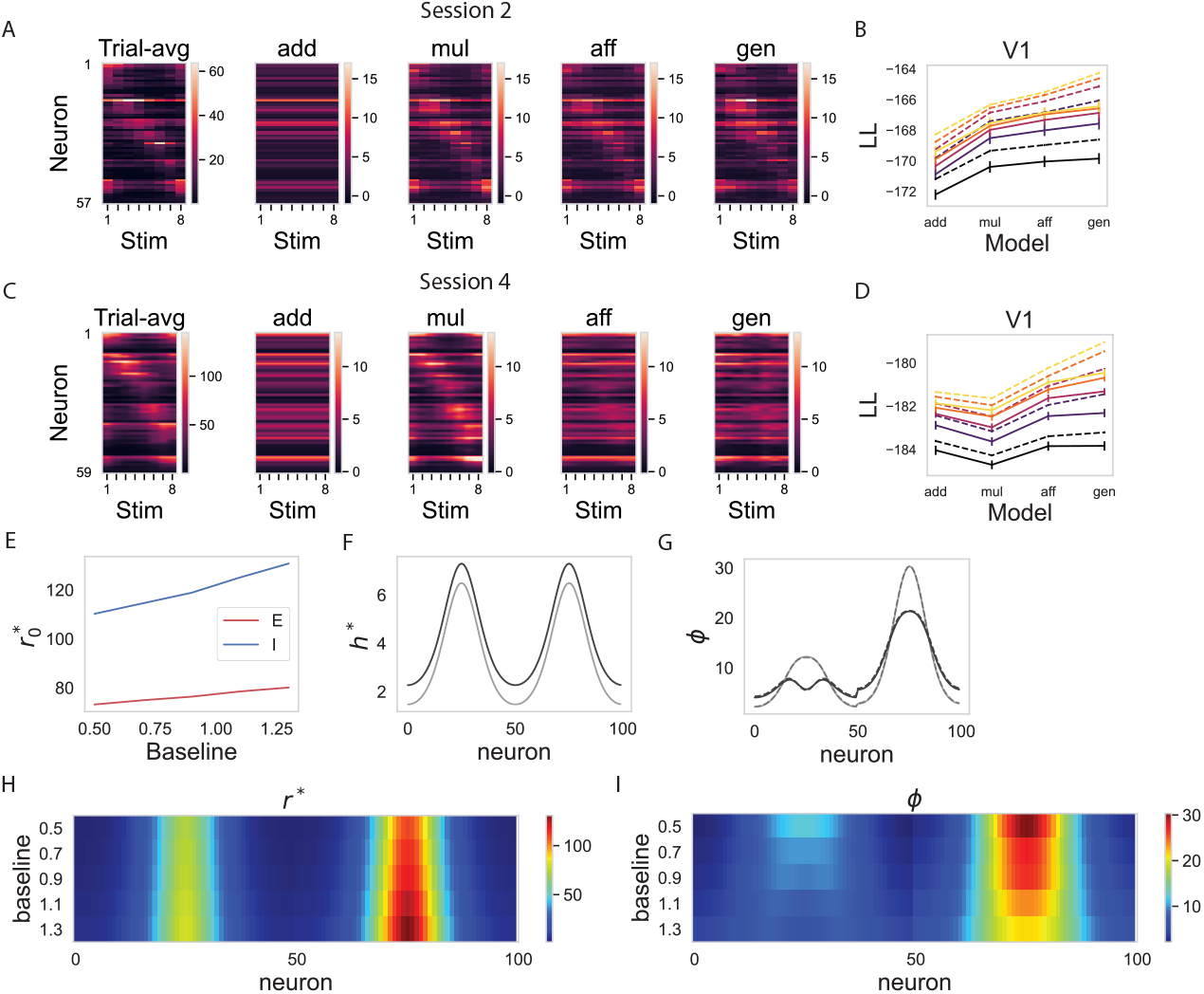
Baseline modulations of the external static inputs account for differences across experimental sessions mechanistically. A-D. Same as in Fig. 3, but for experimental session 2 and session 4. Share variability within V1 for session 2 is multiplicative-like, whereas shared variability within V1 for session 4 is additive-like. E-I Ring model simulation results. E. The simulated time-averaged firing rates of E and I neurons with PO matching the stimulus orientation plotted against stimulus baseline. F. Two example stimuli with low baseline (gray) and high baseline (black). G. The corresponding shared variability amplitudes under the two example stimuli. H. Time-averaged (trial-averaged) activity *r*^*^ increases with increasing the baseline of the external static input. I. Shared variability amplitude *ϕ* for external static inputs with different baseline levels.

Different experimental sessions exhibit qualitative differences in the fitting performances (also reported by (Lin et al., 2015)). For sessions 0, 1 and 2, for both V1 and V2, multiplicative models outperform additive models. However, for V1 data in sessions 3 and 4, additive models outperform multiplicative models (Supplementary Fig. S5, Supplementary Fig. S6A-D). The circuit model can capture the differences between sessions by adjusting a baseline of external static input (Supplementary Fig. S6E-I). With increasing baseline, the shared variability amplitude varies less across orientations, and thus becomes better fit by additive models than multiplicative models.

**Figure S7:**
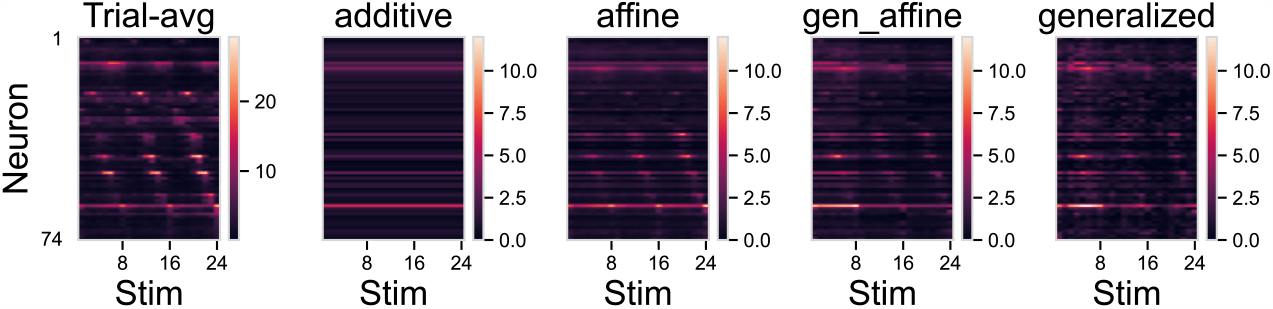
The inferred amplitude of variability shared within V2 from statistical models with 1 component plotted against trial-averaged neuronal responses. The stimuli are drifting gratings with 8 orientations and 3 contrast levels.

**Figure S8:**
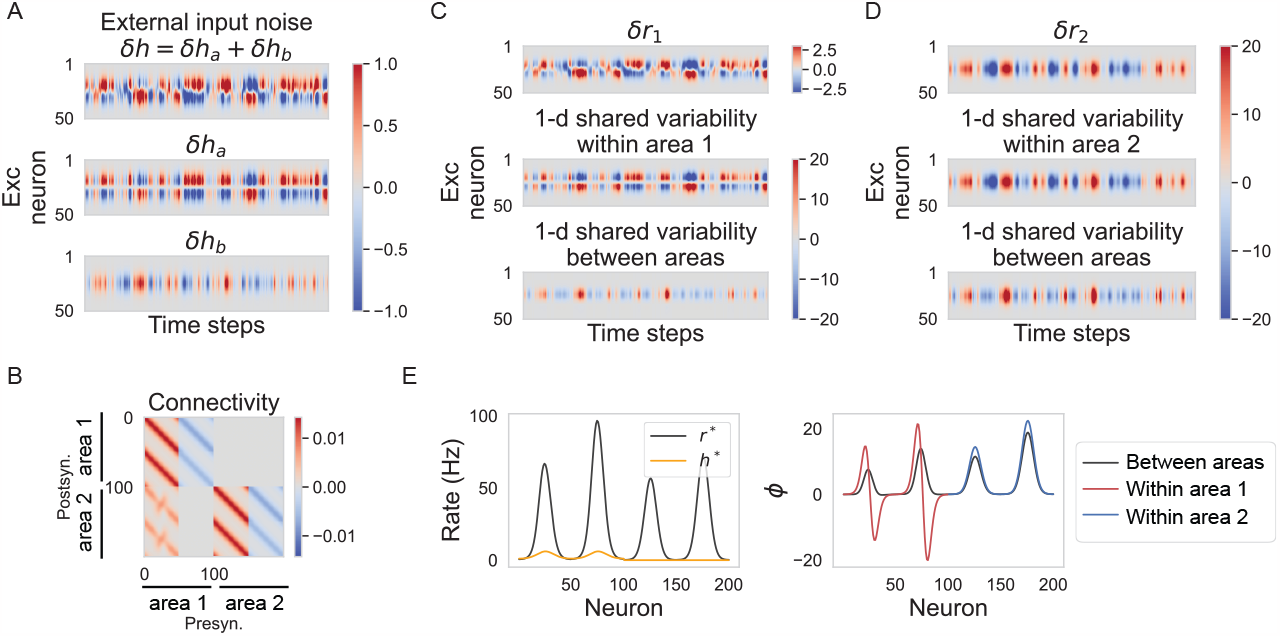
Model simulation shows that it is possible to have different stimulus-dependence for shared variability within and between areas. A. Top panel: external input noise *δh* received by excitatory neurons in area 1. *δh* is 2-dimensional, we plotted each dimension separately (*δh*_*a*_ and *δh*_*b*_) in the bottom two panels. B. Connectivity between neurons in the two areas. Here we set the connectivity between area 1 and area 2 to be nearly orthogonal to residual responses in area 1 caused by *δh*_*a*_. C. From top to bottom panels: residual activity *δr* of the excitatory neurons in area 1; the dominant 1-dimensional variability shared within area 1; the dominant 1-dimensional variability shared between two areas. D. Similar to C. but for area 2. Note that area 2 does not receive any external input noise directly. E. Left: the external static input *h*^*^ and the time-averaged rate responses *r*^*^ of all the simulated neurons in the connected ring model. Right: the amplitudes of the dominant dimension of shared variability. Red: amplitude of variability shared within area 1. Blue: amplitude of variability shared within area 2. Black: amplitude of variability shared between two areas.

**Figure S9:**
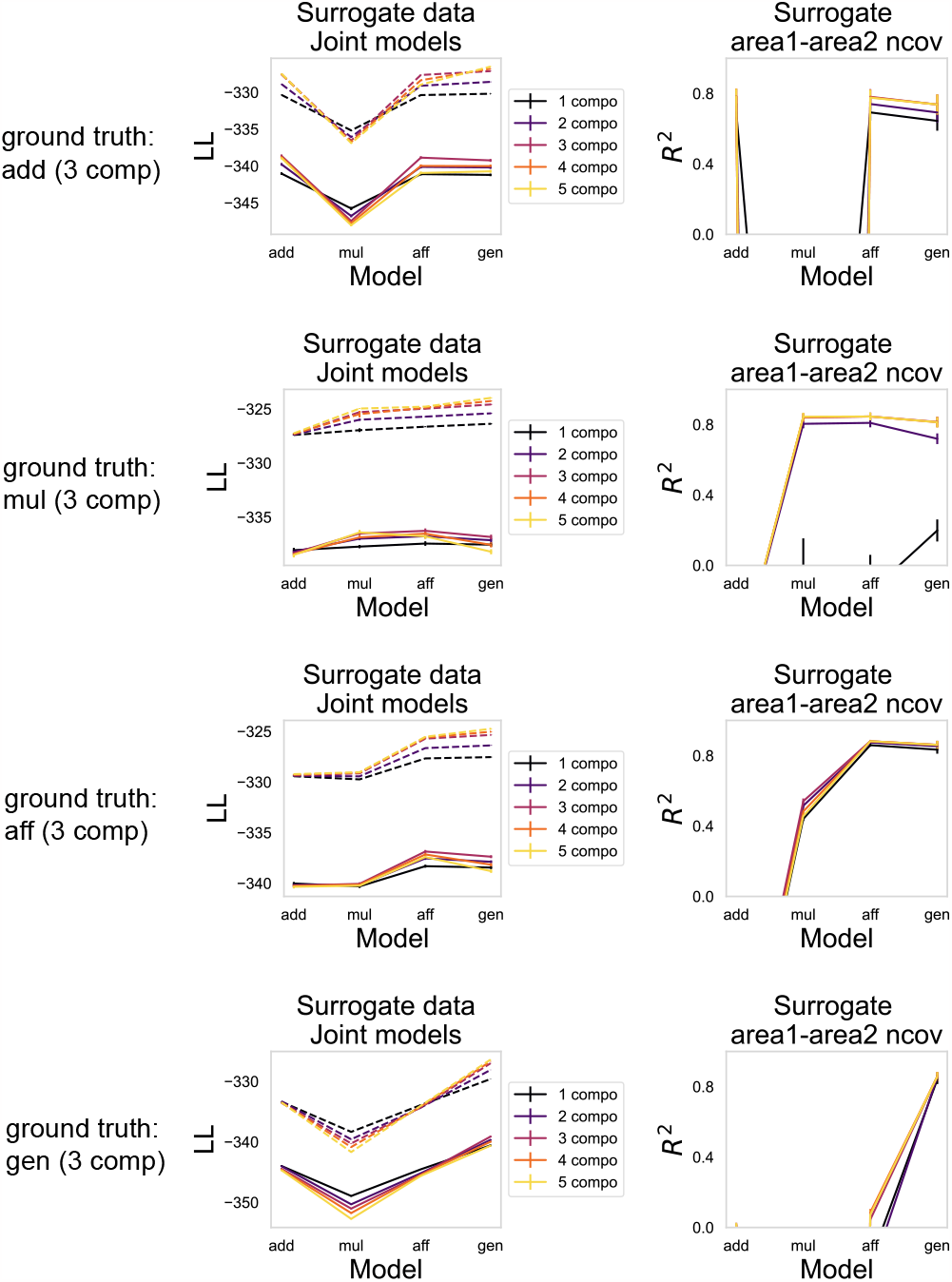
Fitting performances of joint statistical models on surrogate data with known ground truth. We generated surrogate data of 2 brain areas ((40 neurons from area 1 + 60 neurons from area 2) x 400 trials x 8 orientations) with similar statistics to the experimental data we have: 1) each neuron has a bell-shaped orientation tuning curve, with trial averaged response span from 20 to 100 Hz across orientations 2) on each trial, shared variability is simulated as a sample taken from a 3-dimensional standard Gaussian distribution. For a given neuron and stimulus orientation, when the ground truth is “add”/”mul”/”aff”, the amplitude of shared variability follows an additive/multiplicative/affine function of trial-averaged responses; when the ground truth is “gen”, the amplitude of shared variability follows a periodic function with 3 peaks across orientations. 3) On each trial, for a given neuron and stimulus orientation, private variability is simulated as a sample taken from a Gaussian distribution with zero-mean and a variance equal to its trial-averaged response. 4) for each brain area, we simulated an additional 1-dimensional variability shared within the area. Joint statistical models with the 3 components and the correct assumption for the variability shared between areas achieve the best performance with the least number of parameters. Dashed line shows log likelihood of the training set, solid line shows log likelihood of the test set.

Dashed line shows log likelihood of the training set, solid line shows log likelihood of the test set.

**Figure S10:**
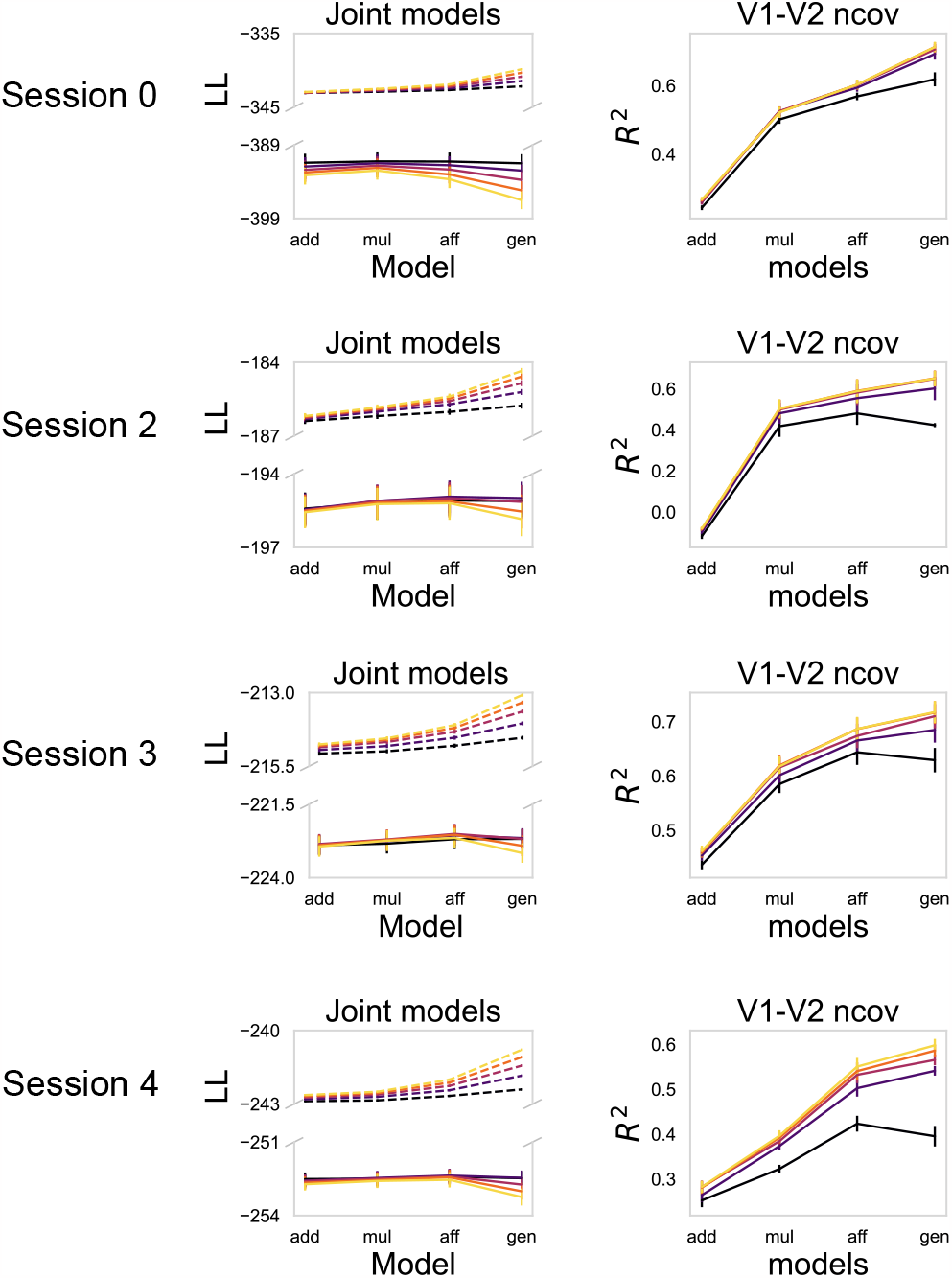
Fitting performances of joint statistical models on V1 and V2 data for session 0,2,3,4 in the dataset 1. Dashed line shows log likelihood of the training set, solid line shows log likelihood of the test set.

**Figure S11:**
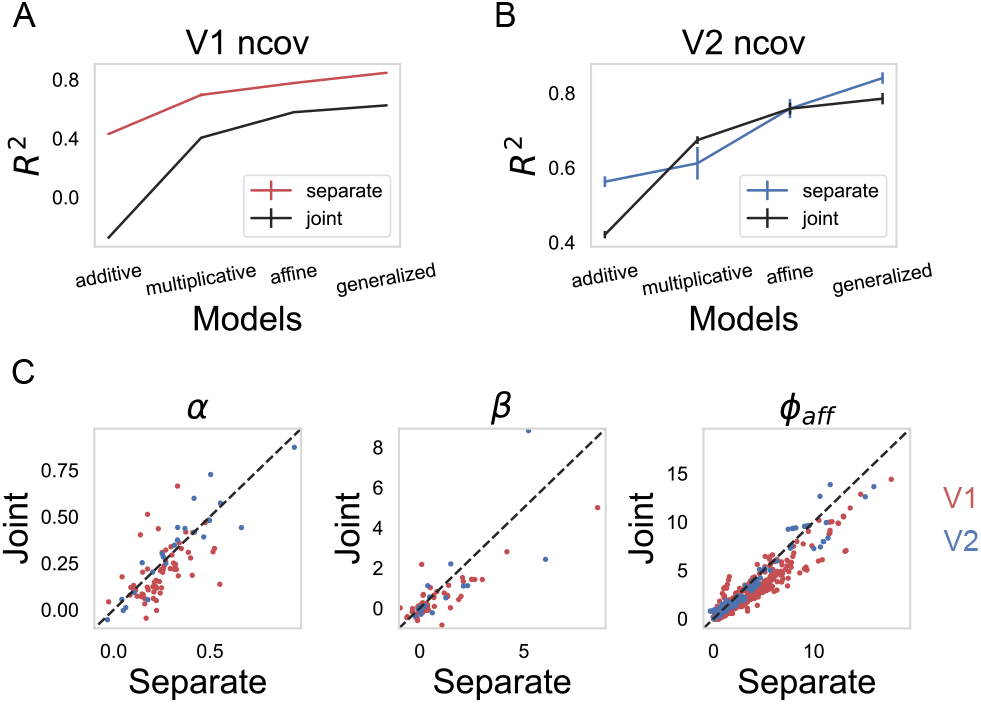
Compare joint statistical models and statistical models fit to V1 and V2 data separately. A., B. The cross-validated *R*^2^ of the noise covariance across stimuli. Error bars show the standard error of 5-fold cross-validation. Colored line denotes the *R*^2^ for the statistical models with 1 component fit to V1 and V2 data separately. Black line denotes the *R*^2^ for the joint statistical models with 1 component. C. The fitted *α, β* and *ϕ* in the affine models and affine joint models with 1 component. Red denotes parameters fit to V1 data, blue denotes parameters fit to V2 data.

**Figure S12:**
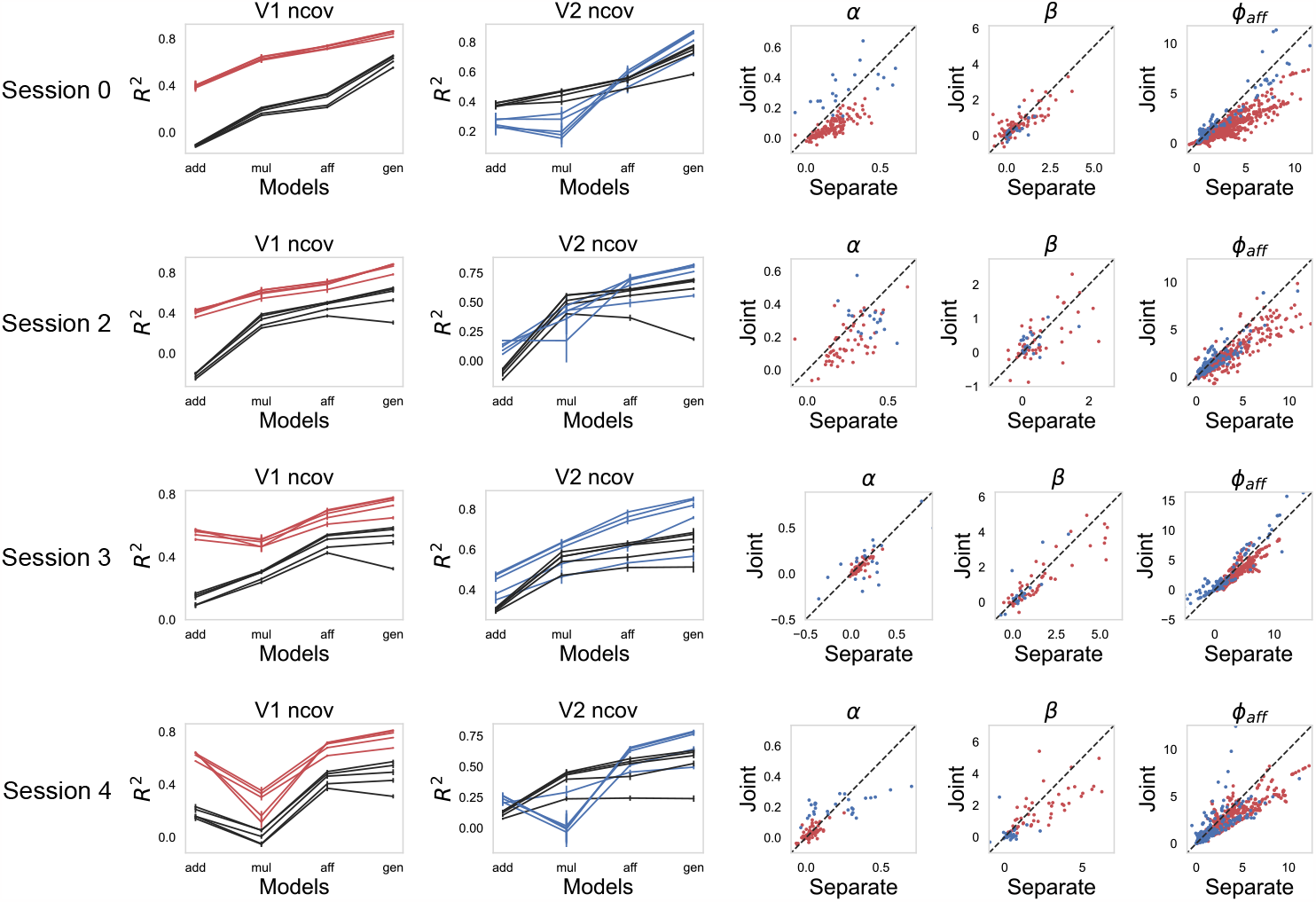
Similar to Supplementary Fig. S11, compare joint statistical models and statistical models fit to V1 and V2 data separately for session 0,2,3,4 in the dataset 1. Different lines with the same color denote *R*^2^ of statistical models with different number of components.

In this note, we analyze how neurons in the ring model respond to external input noise, assuming the noise is small but not necessarily slow. Our goal is to provide an explanation for how shared variability amplitudes depend on the orientations under different contrasts levels.

### S1 Linearized dynamics of the ring model

With N excitatory and N inhibitory neurons sitting in the ring model, their dynamics is described by:

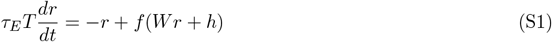

with 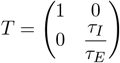, and 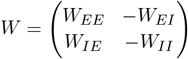.

Consider the fluctuation *δr* around a certain fixed point *r*^*^ = *f* (*Wr*^*^ + *h*^*^) due to external input noise *δh* = *h* − *h*^*^, assume the fluctuation and the input noise to be small, we get the linearized dynamics of *δr*:

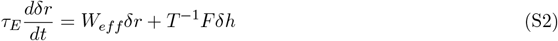

We define the matrix *F* = *diag*(*f*^*′*^(*Wr*^*^ + *h*^*^)), with diagonal elements representing the effective gain, and

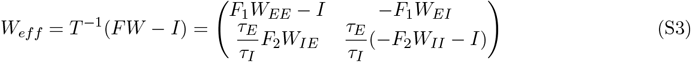

Eq. S2 refers to a 2N-dimensional linear dynamical system. *F*_1_ is diagonal matrix with diagonal elements representing the effective gain of excitatory neurons. Similarly, *F*_2_ is diagonal matrix with diagonal elements representing the effective gain of inhibitory neurons.

Our objective is to develop an intuitive understanding of the behavior of *δr* in response to *δh*. To achieve this, we aim to transform the 2N-dimensional system into a composite of multiple low-dimensional systems. By characterizing the dynamics of these low-dimensional systems, we can extend our understanding to the 2N-dimensional system.

### S2 From 2N-dim linear system to N almost decoupled 2-dim linear system

In the ring model, all the submatrices in *W* are set to be translational-invariant and symmetric (eq. 6). We consider the case where 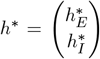 with 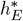 (or 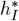) both proportional to a circular Gaussian bump centered around the 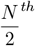 excitatory (or inhibitory) neuron. Accordingly, the diagonal elements of *F*_1_ and *F*_2_ are also proportional to a bump centered around the the 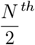 excitatory (or inhibitory) neuron.

From this set up, we can see that

- The submatrices of *W*_*eff*_ are not perfectly symmetric; however, they are nearly symmetric (or nearly normal).
- If we configure the submatrices within *W* to vary solely by scaling and we assume that *F*_1_ and *F*_2_ differ only by scaling as well, the difference between submatrices in *W*_*eff*_ (see eq.S3) would be limited to scaling or the subtraction of multiples of the identity matrix. Therefore, in this case, the submatrices in *W*_*eff*_ will share the same Schur basis, which is denoted as {*e*_*i*_}.

We can write the linearized dynamics described by eq.S2 in the following orthonormal basis comprised of pairs of vectors: 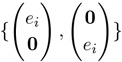, where the set {*e*_*i*_} constitutes the identical Schur basis shared by the submatrices within *W*_*eff*_, and **0** represents an *N* × 1 vector filled with zeros. Intuitively, we refer to 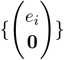 as E modes, as only excitatory neural activities contribute to the dynamics along these modes; similarly, we refer to 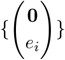 as I modes.

The dynamics of *δr* described in the basis *b*, i.e., 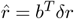, can be expressed as follows:

**Figure S13:**
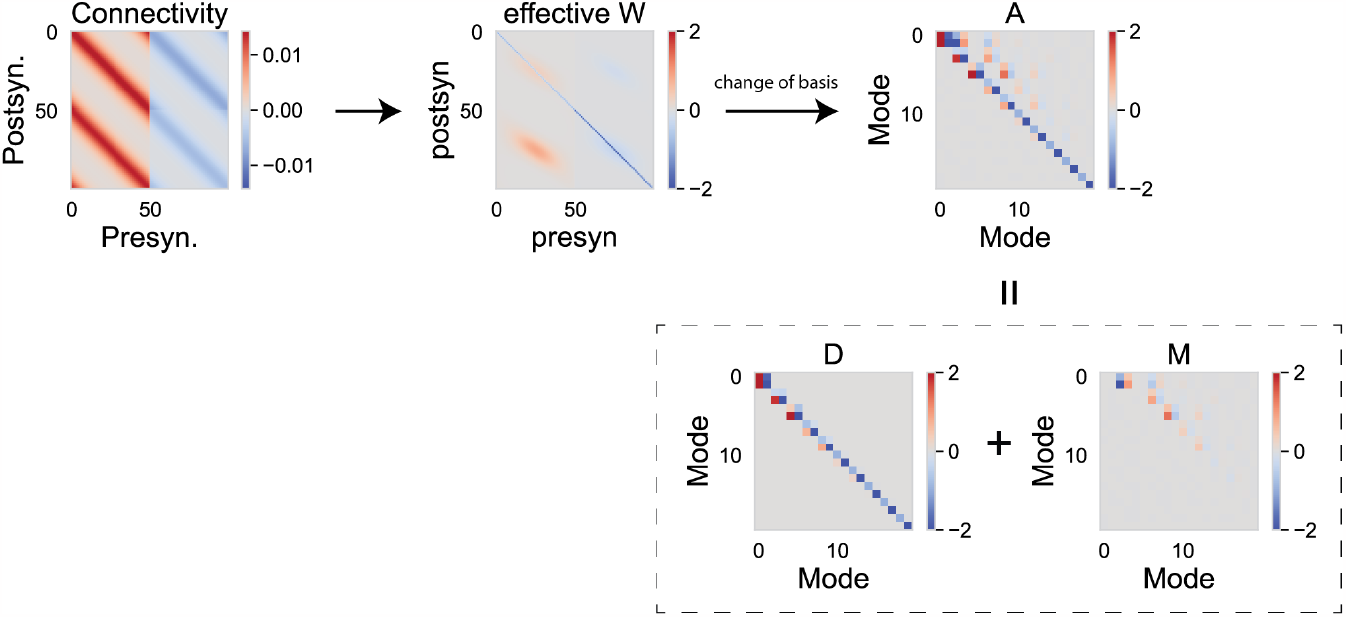
A schematic for illustrating how to get N “almost” decoupled 2-dimensional system from a 2N-dimensional ring circuit model, with a zoomed-in views of matrices *A, D*, and *M*.

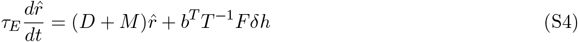

We define *A* = *b*^*T*^ *W*_*eff*_ *b*, where *b* is a 2*N* × 2*N* matrix that represents the previously described orthonormal basis. The columns of *b* consist of E and I modes. We arrange these columns as pairs of E and I modes, and we order these pairs by their associated eigenvalues of the *E* → *I* submatrix of *W*_*eff*_ in descending order.

We express *A* as the sum of two matrices, *D* and *M*. *D* is a block-diagonal matrix comprising N 2 × 2 diagonal blocks, with each element in these 2 × 2 blocks being an eigenvalue of the corresponding submatrix of *W*_*eff*_. *M* contains all the 2 × 2 blocks in the upper triangular entries (Fig. S13). Due to the close-to-normal nature of the submatrices of *W*_*eff*_, the Schur forms of these submatrices approach diagonal matrices. Consequently, most elements within *M* will be zeros.

*A* = *D* + *M* describes the coupling between dynamics along different modes. *D* describes the coupling between E and I mode within the same pair. *M* describes the feedforward coupling between pairs of modes. Because most of the elements within *M* are zeros, the 2N-dimensional linear dynamical system described by eq. S4 is equivalent to N weakly and feedforwardly coupled 2-dimensional linear dynamical systems.

**Figure S14:**
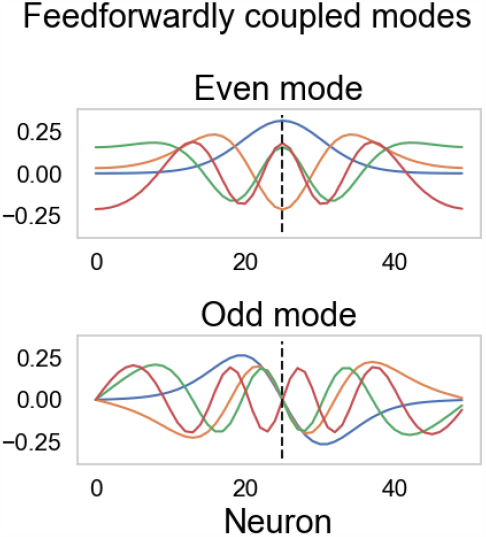
*e*_*i*_ of the leading (lower-order) even and odd modes

#### Dynamics along even modes is decoupled with dynamics along odd modes

We further define “even modes” as basis vectors that involve *e*_*i*_, where *e*_*i*_[*N/*2 − *k*] = *e*_*i*_[*N/*2 + *k*], and “odd modes” as those involving *e*_*i*_, where *e*_*i*_[*N/*2 − *k*] = −*e*_*i*_[*N/*2 + *k*], for any *k* ∈ {0, 1, …, *N/*2} (Fig. S14). Since both *F*_1_ and *F*_2_ are even (i.e., *diag*(*F*)[*N/*2 − *k*] = *diag*(*F*)[*N/*2 + *k*] for any *k* ∈ {0, 1, …, *N/*2}), it follows that for all the submatrices *W*_*sub*_ of *W*_*eff*_, the product *W*_*sub*_*e*_*i*_ has the same parity as *e*_*i*_. Consequently, after the change of basis, even modes and odd modes get decoupled. For instance, 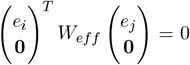 if *e*_*i*_ is even and *e*_*j*_ is odd. In other words, only 2-dimensional systems associated with modes of the same parity are feedforwardly coupled.

#### N decoupled 2-dim systems approximates the 2N-dim system

Due to the sparsity of feedforward couplings between the N 2-dimensional systems, ignoring these feed-forward couplings has minimal impact on the dynamics along each mode, as depicted in Fig. S15. Consequently, we can focus on understanding the dynamics within each 2-dimensional system, which is governed by the corresponding 2 × 2 diagonal blocks within the matrix *D*.

As a reminder, elements in the diagonal blocks of *D* are the eigenvalues of submatrices within *W*_*eff*_. According to the following conditions:

- Each row of *W*_*EE*_ follows a circular Gaussian function, so that *W*_*EE*_ is positive definite
- The largest eigenvalue of *F*_1_*W*_*EE*_ exceeds 1
- Different submatrices of *W* only differ by scaling

We can deduce the signs of elements in the 2 × 2 blocks in *D* transition from 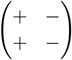 to 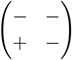 to 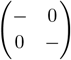, as the indices increase. For the low-order (leading) modes, these block signs indicate non-normal dynamics, implying that we may observe transient increase along E or I mode in their impulse response. In contrast, for the high-order modes, the block signs suggest exponentially decaying dynamics along both E and I mode as their impulse response (Fig. S15).

**Figure S15:**
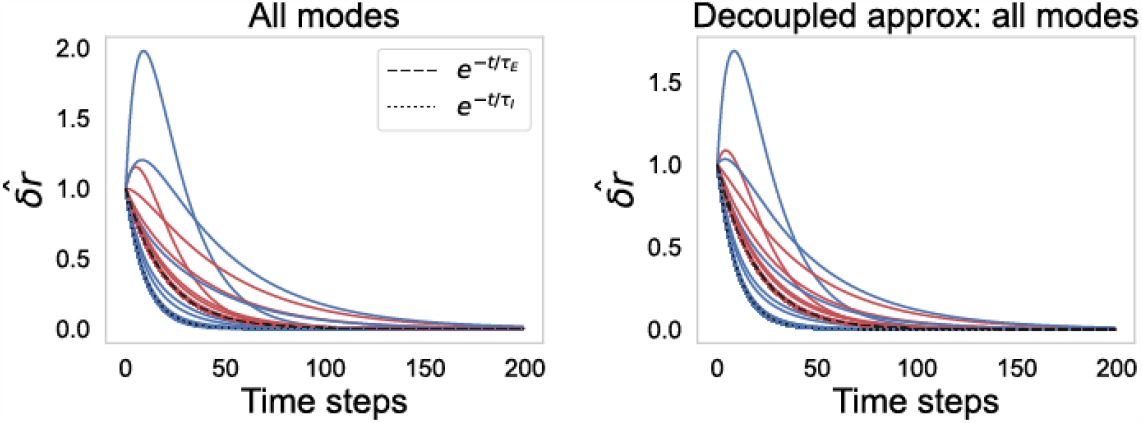
We pick the initial conditions for each mode as 1. And we set *δh* = 0. **Left**, simulated linearized dynamics with *A* = *D* + *M* along all the modes. Blue color denotes dynamics along the Inhibitory modes, red color denotes dynamics along the Excitatory modes. All the higher-order modes have simple dynamics as predicted by the dashed/dotted line (exponential decay). **Right**, the simulated linearized dynamics with *A* ≈ *D* along all the modes. Ignoring the feedforward coupling between the 2-dim linear systems doesn’t change the dynamics qualitatively.

### S3 What is *δr* given *δh*?

The procedure for calculating *δr* from *δh* can be summarized in the following steps. First, we determine the external perturbation received by each mode: *ĥ* = *b*^*T*^ *T*^−1^*Fδh*. Second, we calculate 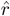 from eq. S4. As suggested by the analysis in the previous section, for the majority of modes *i*, 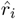 will be the convolution of *ĥ*_*i*_ with an exponentially decaying kernel. In contrast, for some lower-order mode *i*, 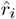 will involve the convolution of *ĥ*_*i*_ with a kernel that exhibits a transient increase. Finally, we calculate *δr* by combining contributions from all the modes 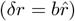.

From this understanding, we infer that *δr* can be multi-dimensional when *δh* is 1-dimensional, and *ĥ* does not perfectly align with any single mode. This arises because 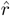 exhibits distinct temporal profiles along different modes.

In the case considered in the main text Fig. 1, where *δh* is slow and identical across neurons, what is 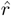 along each mode? Since *δh* is even, for any odd mode *i, ĥ*_*i*_ = 0. Consequently, 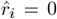 for these odd modes. To further investigate, we examined how the amplitude of 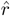 along the leading even E or I mode changes with increasing contrast. We found that the amplitude of the first even E or I mode first increases and then decreases, whereas the amplitude of the second gradually increases and then levels off (Fig. S16). The dominant even mode exhibits a bump-shaped pattern, while the second even mode displays an “M”-shaped pattern (Fig. S14). This observation aligns with the simulation results (main text Fig. 1), where the amplitude of *δr* transitions from a bump-shaped profile to an “M”-shaped profile as contrast levels increase.

**Figure S16:**
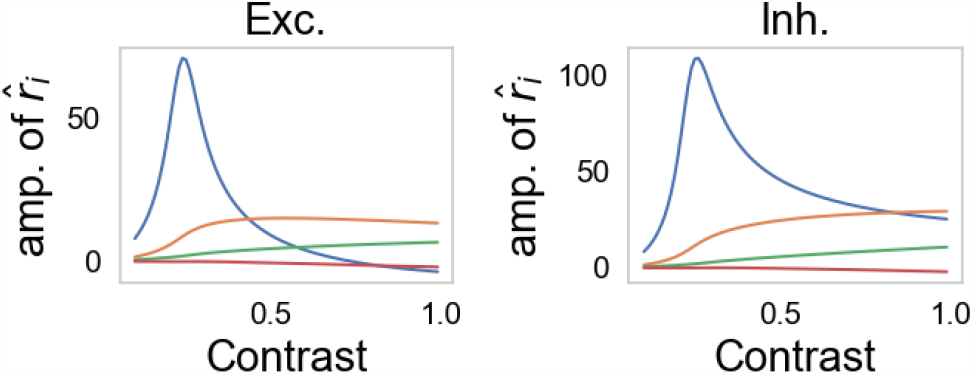
We set *δh*(*t*) to be identical for all the neurons and slow enough to be treated as a constant. **Left**: the amplitude of 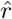 along the leading even E modes, **Right**: the amplitude of 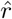 along the leading even I modes. The color code of different even modes are the same as the top panel in Fig. S14

To understand why the amplitudes of the pair of even E and I mode change in this manner, it suffices to examine the behavior of the corresponding 2-dimensional linear system. Because this system is nearly decoupled from the other 2-dimensional linear systems. In essence, we only need to look at how *ĥ* along the E and I mode of interest and the corresponding 2 × 2 block in *D* behave.

Here, we analyzed the dynamics in the 2-dimensional linear system corresponding to the 1st and the 2nd even modes (Fig. S17). As contrast level increases, both *F*_1_ and *F*_2_ increase, leading to a proportional rise in the effective perturbation (*ĥi*).(To intuiti vely assess the behavior of the *i*^*th*^ 2 × 2 block in matrix *D* (*D*_*i*_), we compute its Schur form: 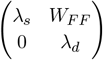. Notably, the *D*_*i*_ matrices corresponding to the 1st and 2nd even modes consistently exhibit sign patterns of 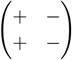 across all contrast levels. Consequently, their Schur vectors are a “sum vector” where both the E mode and the I mode have the same sign and a “difference vector” where the E mode and the I mode have different signs. The eigenvalues *λ*_*s*_ and *λ*_*d*_ characterize the decay rates of dynamics along these sum and difference Schur vectors, with both eigenvalues being negative. Moreover, *W*_*FF*_ represents the feedforward weight that couples the dynamics of these Schur vectors. As derived in (Hennequin et al 2018), the total variance of 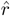 is proportional to 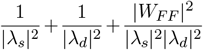. Consequently, the amplitude of 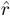 increases as |*λ*| decreases and |*W*_*F F*_ | increases.

To elucidate the contrast-dependent behavior of 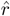 along the 1st even E and I mode (depicted as the blue curve in Fig. S16 and the upper panel in Fig. S17), we observe that the transient decrease in |*λ*_*s*_|, along with the increase in |*W*_*F F*_ | and *ĥ*, collectively contribute to the transient elevation of 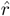. Subsequently, the escalation of |*λ*| accounts for the decline in 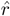 at high contrast levels.

In the context of the contrast-dependent behavior of 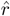 along the 2nd even E and I mode (illustrated by the orange curve in Fig. S16 and the lower panel in Fig. S17), we observe that the transient reduction in |*λ*_*s*_|, along with the increase in |*W*_*F F*_ | and *ĥ*, similarly contribute to the transient increase in 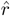.” Furthermore, the saturation of |*λ*| plays a pivotal role in the saturation of 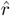 at high contrast levels.

**Figure S17:**
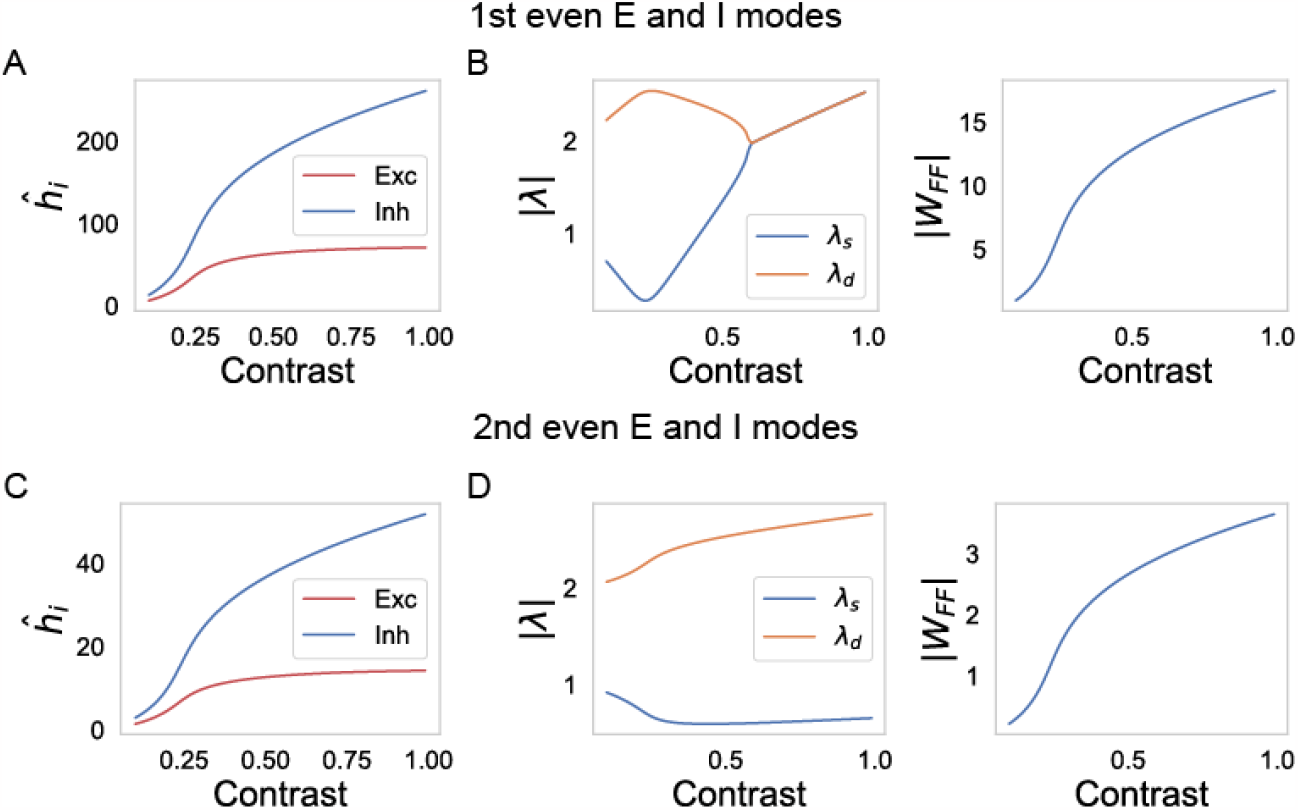
A. The contrast-dependence of *ĥ* along the 1st even E and I mode. B. The contrast-dependence of |*λ*_*s*_|, |*λ*_*d*_| and |*W*_*F F*_ | in the Schur form of the 2 × 2 block in *D* corresponding to the 1st even modes. C, D is similar to A, B but for the 2nd even E and I modes.

